# Small Molecule Modulation of the Archetypal UbiB protein COQ8

**DOI:** 10.1101/2022.03.22.485346

**Authors:** Nathan H. Murray, Adam Lewis, Christopher R. M. Asquith, Juan P. Rincon Pabon, Zixiang Fang, Naomi Ptak, Robert W. Smith, James D. Vasta, Chad A. Zimprich, Cesear R. Corona, Matthew B. Robers, Craig A. Bingman, Michael L. Gross, Katherine Henzler-Wildman, David J. Pagliarini

## Abstract

Small molecule tools have enabled mechanistic investigations and therapeutic targeting of the protein kinase-like (PKL) superfamily. However, such tools are still lacking for many PKL members, including the highly conserved and disease-related UbiB family. Here, we sought to develop and characterize inhibitor and activator molecules for the archetypal UbiB member, COQ8, whose function is essential for coenzyme Q (CoQ) biosynthesis. Guided by crystallography, activity assays, and cellular CoQ measurements, we repurposed the 4-anilinoquinoline scaffold to selectively inhibit human COQ8A in cells. Second, using ^1^H-^13^C HMQC NMR and hydrogen-deuterium exchange mass spectrometry, we reveal that the CoQ precursor mimetic, 2-propylphenol (2-PP), modulates the quintessential UbiB KxGQ domain to increase COQ8A nucleotide affinity and ATPase activity. Our newfound chemical tools promise to lend new mechanistic insights into the activities of these widespread and understudied proteins and to offer potential therapeutic strategies for human diseases connected to their dysfunction.

## Introduction

The protein kinase-like (PKL) superfamily is an expansive set of more than 500 human proteins that perform critical roles in numerous biological processes^1^. Our understanding of these roles has been greatly advanced by the development of molecular tools capable of manipulating PKL activity^2^. Small molecule inhibition of PKL members has also emerged as a remarkably effective therapeutic strategy, with more than 70 new drugs approved in the past 20 years^3, 4^. However, such tools and drugs are still lacking for many PKL members. Approximately 150 human PKL members are still considered to be understudied, including a large number of atypical kinases and pseudokinases^5, 6^. Efforts to expand the repertoire of molecular tools for these proteins promise to further accelerate our understanding of their diverse functionality and expand the possibilities of manipulating their activities for therapeutic gain^7^.

The UbiB proteins are a subset of atypical and understudied PKL family members. These proteins are conserved throughout all domains of life^8^ and, in eukaryotes, are found in cellular compartments responsible for the synthesis and distribution of prenylated metabolites^9–11^. The most well-studied UbiB members are the *Saccharomyces cerevisiae* protein Coq8p and its corresponding human homologs, COQ8A and COQ8B (collectively referred to here as COQ8). COQ8A and COQ8B each have well established connections to human disease, with inactivating mutations resulting in autosomal recessive cerebellar ataxia 2^12–15^ and steroid-resistant nephrotic syndrome^16^, respectively. Early investigations identified COQ8 proteins as auxiliary factors required for the biosynthesis of coenzyme Q (CoQ)^17, 18^, a redox-active prenyl lipid essential for biological processes including cellular respiration, pyrimidine biosynthesis, and cellular antioxidation^19^. COQ8 was initially proposed to be a protein kinase based on sequence homology, but structure/function studies of COQ8A revealed conserved features poised to prevent canonical protein kinase activity^20^. Instead, COQ8 may use its ATPase activity to maintain the integrity of a CoQ biosynthetic complex and to access hydrophobic CoQ intermediates embedded in the mitochondrial inner membrane^21^; however, this model requires further validation. Similar to other PKL members, the development of new molecular tools to manipulate COQ8 function could help elucidate its precise molecular function and establish it as a therapeutic target for human disease.

Previous efforts have set a foundation for the development of small molecular activators and inhibitors of COQ8^22^. We recently developed a covalent inhibitor against an engineered yeast Coq8p double mutant (M303C, V202C), but which lacked efficacy against the WT protein^21^. Furthermore, using NMR and activity screens, we identified a set of CoQ headgroup-like phenolic compounds, including 2-propylphenol (2-PP) and 2-allylphenol (2-AP), that enhance COQ8 ATPase activity^21^; however, the binding site and mechanism of activation remain unclear. To our knowledge, no activators or inhibitors of WT COQ8 proteins have been validated in a biological context. Here, to address these limitations, we repurposed a 4-anilinoquinoline scaffold to create a custom endogenous inhibitor of COQ8A, and helped reveal the mechanism by which 2-PP activates COQ8A. Collectively, this work expands the PKL small-molecule toolkit and offers new resources to investigate the understudied UbiB family and its connection to human disease.

## Results

### 4-Anilinoquin(az)olines bind and inhibit COQ8A *in vitro*

Functional investigations of COQ8 have been hindered by the lack of established inhibitors. To identify lead inhibitor compounds, we searched published kinase screening data and found COQ8A as a promising secondary target for 4-anilinoquinolines^23, 24^. To further explore this scaffold, we tested the ability of 35 distinct 4-anilinoquin(az)oline compounds to bind COQ8A by using differential scanning fluorimetry (DSF) (Fig. 1a, 1b). Twenty-five of these compounds increased COQ8A melting temperature significantly more than our weak-binding control, erlotinib. The strongest stabilizers featured methoxy groups on the aniline ring at the *para*-and both *meta*-positions and electronegative atoms at positions 6 or 7, suggesting important molecular features for COQ8A binding. We further tested the top eight compounds in an *in vitro* COQ8A ATPase activity assay in the presence of 2-PP and Triton X-100, compounds we previously found to enhance COQ8A activity (Fig. 1c, Extended Data Fig. 1a). All tested compounds inhibited COQ8A^NΔ250^ (which includes the full PKL domain and lacks only the mitochondrial targeting sequence and single-pass transmembrane domain)^20^ with IC_50_ values ranging from 0.47 to 1.7 μM. By DSF, these compounds possess *K_d,app_* values ranging from 7.6 to 27 μM (Fig. 1d, Extended Data Fig. 1b).

**Fig. 1.**
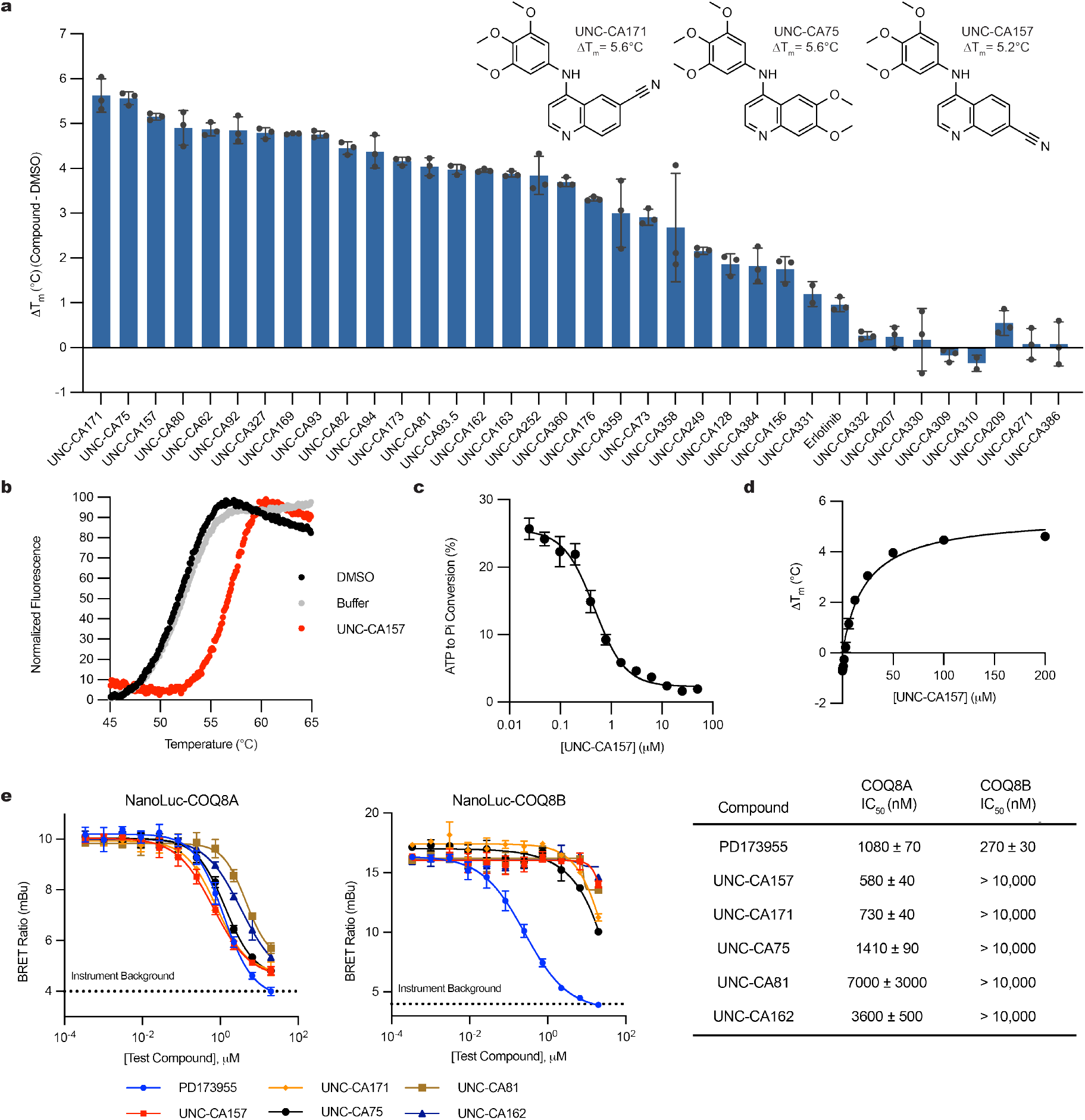
Characterization of *in vitro* COQ8A inhibitors. **a**, Differential scanning fluorimetry (DSF) screen of 4-anilinoquinoline compounds (n=3 ± SD). **b**, DSF traces of COQ8A^NΔ250^ with UNC-CA157 compared to controls. **c**, Inhibition of COQ8A^NΔ250^ ATPase activity by UNC-CA157 (n=3 ± SD). **d**, UNC-CA157 binding curve, as determined by DSF (n=3 ± SD). **e**, Live cell target engagement for COQ8A (left) and COQ8B (right) assessed in HEK293 cells using a NanoBRET approach (n=3 ± SD representative of two independent experiments). PD173955 served as a control.

To test the ability of these compounds to engage COQ8A and COQ8B in live HEK293 cells, we developed a bioluminescence resonance energy transfer (BRET) assay based on the NanoLuc luciferase technology^25^. This NanoBRET target engagement assay includes COQ8A or COQ8B fused to an N-terminal NanoLuc and a COQ8 BRET probe. In the absence of inhibitor the COQ8 BRET probe and fusion protein generated a detectable BRET signal. Competitive displacement of the COQ8 BRET probe from the ATP binding site by our inhibitor series resulted in a dose-dependent loss of BRET signal from which we determined IC_50_ values. With the caveat that N-terminal tagging likely prevents COQ8 mitochondrial localization, 4-anilinoquinolines showed a clear preference for COQ8A over COQ8B and UNC-CA157 showed the lowest IC_50_ against COQ8A at 580 nM. From these data, we selected UNC-CA157 as the most promising lead candidate for further development.

### UNC-CA157 slightly inhibits CoQ production in COQ8B^KO^ cells

We next set up a system to test the efficacy of our inhibitor in human HAP1 cells. In humans, COQ8A and COQ8B are paralogs that likely possess redundant functions in the CoQ biosynthesis pathway and that exhibit some tissue specificity^14, 16, 26^. In a recent study (Rensvold et al. 2022, accepted), we determined that both paralogs are active in HAP1 cells, and that COQ8B is the more dominant version whose disruption causes the accumulation of the early CoQ precursor, polyprenyl hydroxybenzoate (PPHB_10_) (Fig. 2a). We performed mass spectrometry measurements and confirmed that the COQ8A^KO^ HAP1 cells had no significant change in cellular CoQ_10_ levels, whereas the COQ8B^KO^ cells had slight CoQ_10_ loss (Fig. 2b). Importantly, the double knockout line (COQ8A/B^DKO^) exhibited marked loss of CoQ_10_, demonstrating that each paralog can largely support the pathway in the absence of the other. The concomitant buildup of PPHB_10_ followed the same phenotypic pattern, with the largest increase seen in the COQ8A/B^DKO^ cells (Fig. 2c). We further measured the impact of our COQ8 protein knockouts on cellular respiration by using seahorse respirometry and again found that disrupting both COQ8 paralogs produced the strongest effect on basal and uncoupled oxygen consumption rates (OCR) (Fig. 2d). Together, these data demonstrate that both COQ8A and COQ8B are active in HAP1 cells, with COQ8B serving as the predominant paralog, thereby presenting a system to test the efficacy and specificity of our inhibitor.

**Fig. 2.**
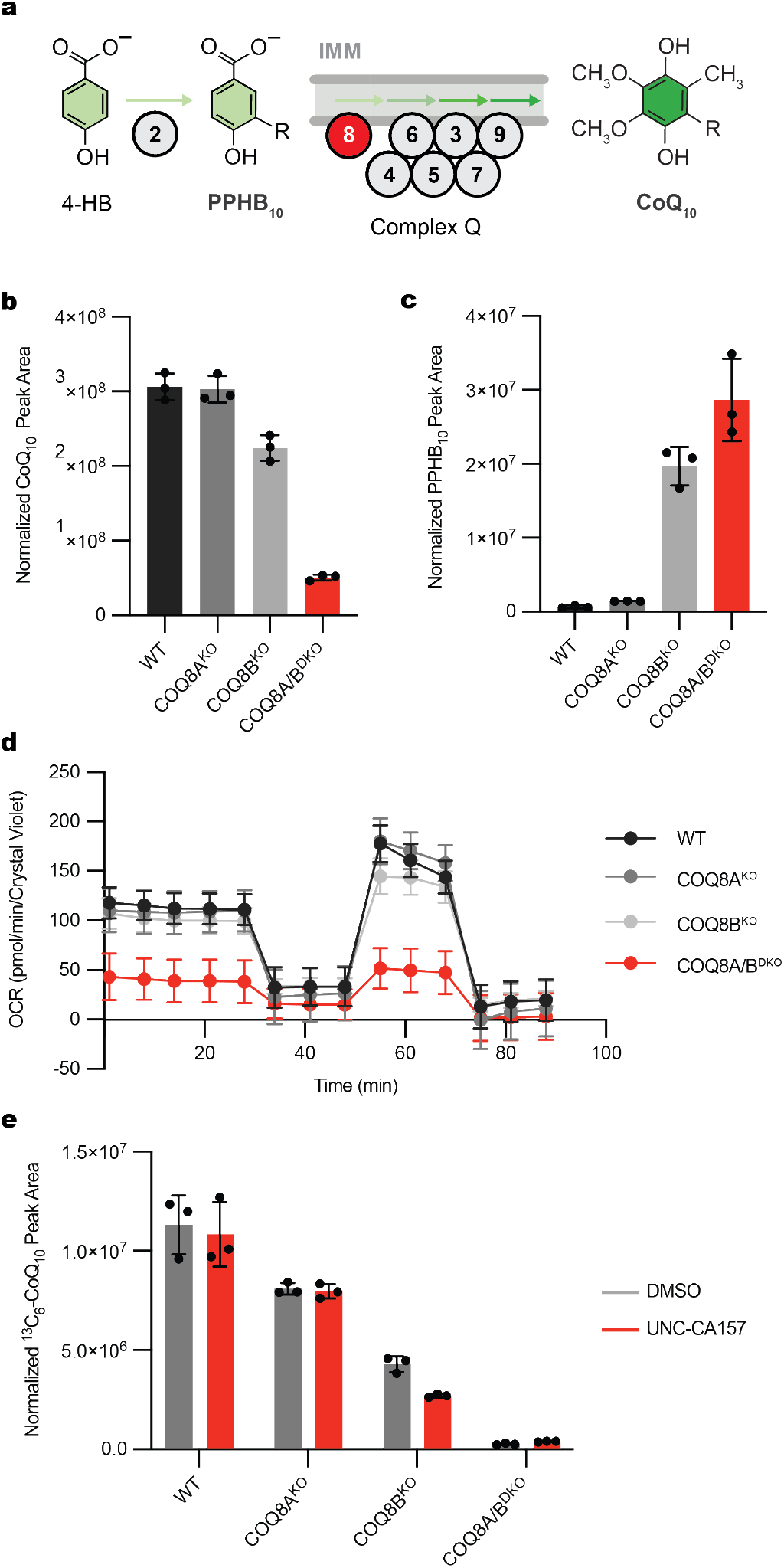
UNC-CA157 decreases *de novo* CoQ production in COQ8B^KO^ HAP1 cells. **a**, Abbreviated schematic of the CoQ biosynthesis pathway. COQ8 proteins are essential for converting PPHB_10_ to mature CoQ_10_. **b**, Total CoQ_10_ levels in WT, COQ8A^KO^, COQ8B^KO^, and COQ8A/B^DKO^ HAP1 cells as determined by LC-MS (n=3 ± SD). **c**, Total PPHB_10_ levels in WT, COQ8A^KO^, COQ8B^KO^, and COQ8A/B^DKO^ HAP1 cells as determined by LC-MS (n=3 ± SD). **d**, Oxygen-consumption rates from a Seahorse mitochondria stress test of WT, COQ8A^KO^, COQ8B^KO^, and COQ8A/B^DKO^ HAP1 cells (n=11-12 ± SD). **e**, ^13^C_6_-CoQ_10_ (*de novo* synthesized CoQ_10_) levels in WT, COQ8A^KO^, COQ8B^KO^, and COQ8A/B^DKO^ HAP1 cells after treatment with 10 μM ^13^C_6_-4-HB and either DMSO or 20 μM UNC-CA157 (n=3 ± SD).

To begin, we measured the impact of UNC-CA157 on *de novo* CoQ production. We treated WT, COQ8A^KO^, COQ8B^KO^, and COQ8A/B^DKO^ HAP1 cells with either 20 μM UNC-CA157 or matched DMSO and monitored the incorporation of the CoQ precursor ^13^C_6_-4-hydrobenzoate (^13^C_6_-4-HB), into CoQ biosynthesis via liquid chromatography-mass spectrometry (LC-MS) (Extended Data Fig. 2a). Incorporation of the labeled 4-HB precursor followed the same phenotypic pattern observed in our cellular CoQ measurements, with single COQ8 knockouts showing decreased ^13^C_6_-CoQ_10_, and a further depletion when both paralogs were disrupted (Fig. 2e). Treatment with UNC-CA157 resulted in a significant decrease in ^13^C_6_-CoQ_10_ production only in the COQ8B^KO^, where cells are solely reliant on the COQ8A paralog for CoQ biosynthesis. No increase in ^13^C_6_-PPHB_10_ or unlabeled PPHB_10_ was seen, suggestive of a relatively weak inhibition (Extended Data Fig. 2b,d). Unsurprisingly, this short incubation with UNC-CA157 had no effect on unlabeled CoQ_10_ levels, given the long half-life of CoQ_10_ (Extended Data Fig. 2c). This was the first demonstration of COQ8A inhibition in a biological context and set the foundation for further probe development around this scaffold.

### UNC-CA157 is a type I kinase inhibitor

To further investigate the inhibitor-protein interaction, we purified COQ8A^NΔ254^, crystallized it in the presence of UNC-CA157, and solved two inhibitor-bound crystal structures (PDB ID=7UDP and 7UDQ) (Fig. 3a, Extended Data Table 1). The structures revealed that UNC-CA157 is a type I kinase inhibitor^27^ that binds to the COQ8A ATP pocket with the αC helix in the ‘in’ conformation and the DFG motif in an ‘intermediate’ conformation (Fig. 3b). Superimposing the UNC-CA157-bound structure with our previous nucleotide-bound structure (PDB=5i35)^15^ reveals that the quinoline ring from UNC-CA157 overlays with the adenosine ring from AMP-PNP, further suggesting direct competition with ATP as the mechanism of inhibition (Fig. 3c). The UNC-CA157-bound structures largely mimic the nucleotide-bound state of COQ8A, resulting in similar modifications to the KxGQ domain—the unique and invariant domain of the UbiB family—when compared to the apo form^15^.

**Fig. 3.**
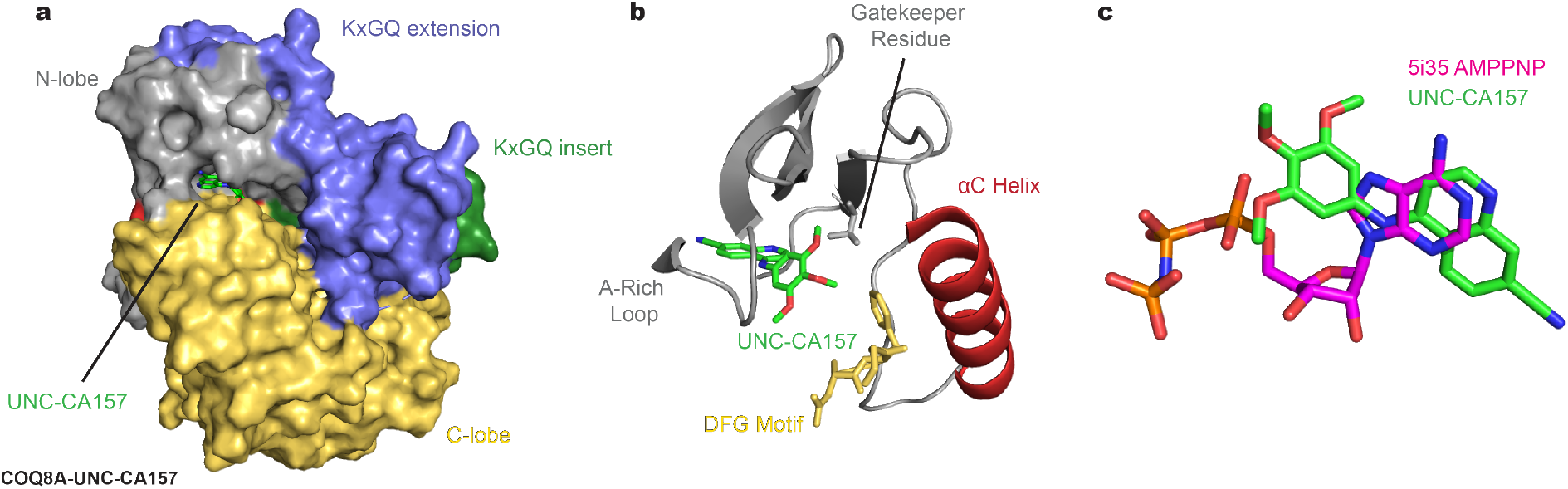
COQ8A-UNC-CA157 crystal structure reveals type I kinase inhibition. **a**, Surface representation of UNC-CA157-bound COQ8A^NΔ254^. **b**, UNC-CA157 bound to the protein active site reveals type I kinase inhibition. **c**, Overlay of AMPPNP (PDB=5i35) and UNC-CA157 in the COQ8A active site.

COQ8A and COQ8B exhibit high sequence similarity; however, the kinase screening data from Asquith et al.^23^ and our cellular data above indicate that UNC-CA157 possesses specificity for COQ8A. To investigate what enables this specificity, we mutated select COQ8A^NΔ250^ active site residues that are in close proximity to UNC-CA157 in our crystal structures and purified the corresponding protein (Extended Data Fig. 3a). These mutations yielded variable effects, with some leading to slightly enhanced inhibitor binding and three causing near complete loss of binding as assessed by DSF (Extended Data Fig. 3b). Among these three, the F495A mutant stood out as particularly noteworthy given that this residue is non-conserved among the human and yeast UbiB proteins, with all but COQ8A having leucine at this position (Extended Data Fig. 3b,c). We further tested this residue by purifying human COQ8A^NΔ250^ WT and F495L (converting phenylalanine to the typical residue found at this position within the family), along with *S. cerevisiae* Coq8p^NΔ41^ WT and L353F (introducing a phenylalanine into a homolog that lacks it). Substitution of this phenylalanine in the human protein again diminished binding and, notably, introduction of this residue into the yeast protein more than doubled inhibitor-induced thermal stability (Extended Data Fig. 3e). Additionally, we tested UNC-CA157 inhibition of human WT and F495L COQ8A^NΔ250^ ATPase activity and observed a greater than 10-fold increase in IC_50_ for the F495L mutant (Extended Data Fig. 3f). Overall, these results identify a COQ8A-specific phenylalanine required for UNC-CA157 potency.

### TPP-UNC-CA157 specifically targets COQ8A

A further motivation for crystallizing UNC-CA157-bound COQ8A was to inform medicinal chemistry efforts toward improving inhibitor efficacy. COQ8A is the only known PKL protein that both resides in mitochondria and binds to 4-anilinoquin(az)olines^23^. Therefore, targeting UNC-CA157 to the mitochondrial matrix is a reasonable strategy to improve our compound specificity. The most common strategy for targeting a molecule to the mitochondrial matrix is to amend it with a lipophilic cation, such as triphenylphosphonium (TPP)^28^. Based on membrane potential and cellular conditions, greater than 1000-fold enrichments in mitochondria can be achieved using this strategy^29^. Our structure reveals that the cyano group of UNC-CA157 is positioned in the solvent-exposed portion of the ATP binding site and is, therefore, well positioned for further modification (Fig. 4a). We derivatized UNC-CA157 by replacing the cyano group with a carboxylate ester followed by a 9-carbon linker and TPP (Fig. 4b)^30^. This new compound, TPP-UNC-CA157, also inhibited COQ8A^NΔ250^ ATPase activity *in vitro*, albeit with a lower potency than UNC-CA157 alone (Fig. 4c). Despite this, we reasoned that the enrichment of this compound in the matrix should overcome decreased potency and simultaneously improve its specificity by increasing its local cellular concentration.

**Fig. 4.**
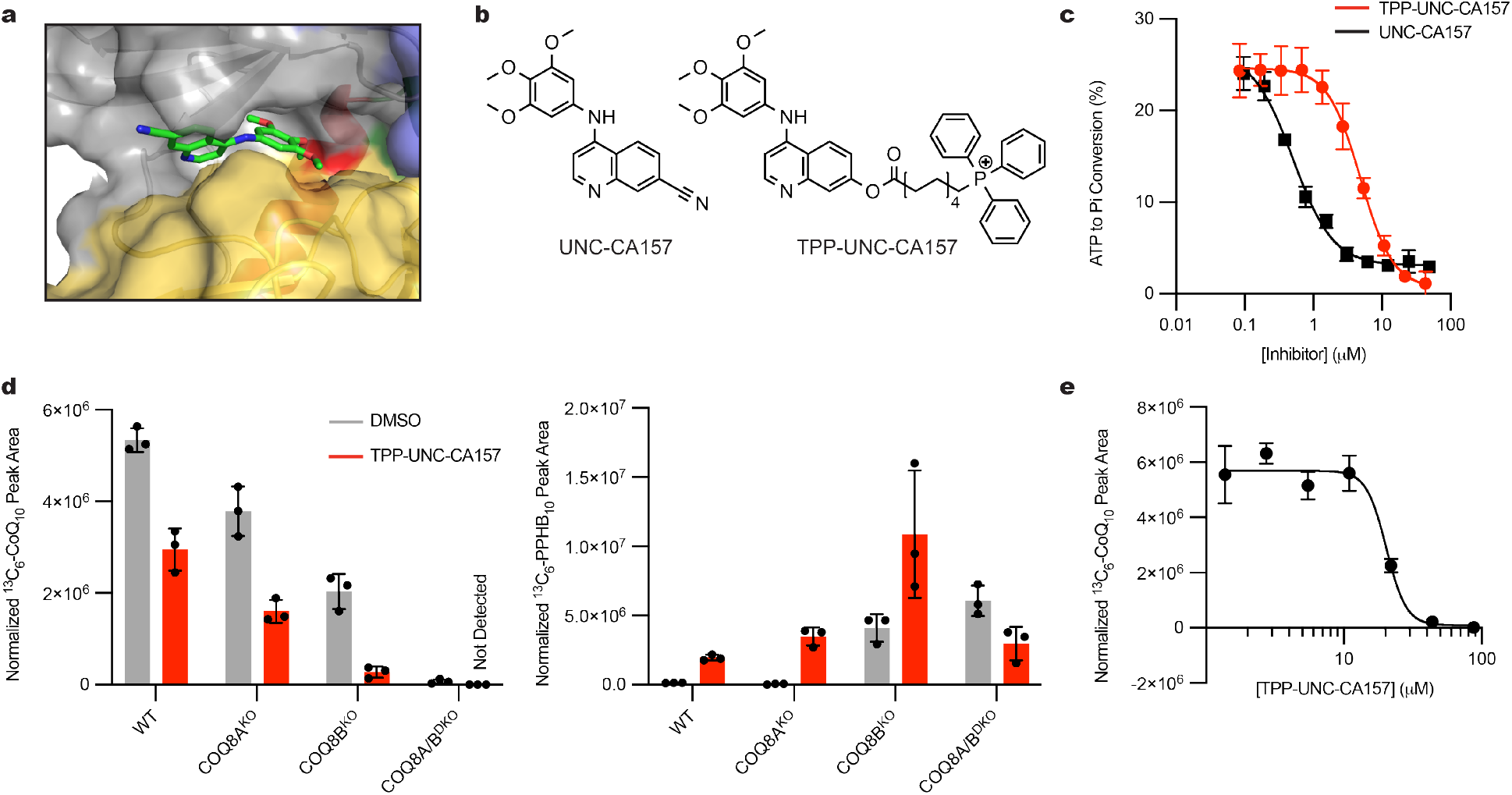
Mitochondrial targeting of TPP-UNC-CA157 increases cellular efficacy in HAP1 cells. **a**, UNC-CA157 positioning in the COQ8A active site revealed an aqueous-exposed cyano group. **b**, The addition of a triphenylphosphonium group to UNC-CA157 for mitochondrial targeting. **c**, Inhibition of *in vitro* COQ8A^NΔ250^ ATPase activity by UNC-CA157 and TPP-UNC-CA157 (n=3 ± SD). **d**, ^13^C_6_-CoQ_10_ (*de novo* synthesized CoQ_10_) and ^13^C_6_-PPHB_10_ (*de novo* synthesized PPHB_10_) levels in WT, COQ8A^KO^, COQ8B^KO^, and COQ8A/B^DKO^ HAP1 cells after treatment with 10 μM ^13^C_6_-4-HB and either DMSO or 17.6 μM TPP-UNC-CA157 (n=3 ± SD). **e**, ^13^C_6_-CoQ_10_ levels in WT HAP1 cells after treatment with 10 μM ^13^C_6_-4-HB and 1.37-87.8 μM TPP-UNC-CA157 (n=3 ± SD).

Treatment of HAP1 cells with TPP-UNC-CA157 resulted in a robust reduction in *de novo* CoQ production, with significant decreases observed in WT, COQ8A^KO^, and COQ8B^KO^ cell lines (Fig. 4d). TPP-UNC-CA157 treatment also led to a reciprocal increase in ^13^C_6_-PPHB_10_ levels, confirming specific disruption of the headgroup modification portion of the CoQ biosynthetic pathway (Fig. 4d). This compound likewise appears to be more efficacious against COQ8A than COQ8B, with the strongest phenotype observed in the COQ8B^KO^ cells. We again observed no changes in unlabeled CoQ_10_ but, owing to the low endogenous levels, a buildup of unlabeled PPHB_10_ was observed (Extended Data Fig. 4a). To determine the potency of our newfound inhibitor, we treated WT HAP1 cells with various concentrations of TPP-UNC-CA157 and were able to abolish completely *de novo* CoQ production in a dose-dependent manner (Fig. 4e). This compound is immediately useful as a tool for interrogating the CoQ biosynthesis pathway and, more broadly, demonstrates the promise of leveraging organellar targeting moieties for selective inhibition of mitochondrial processes. Nonetheless, owing to observed toxicity of TPP-UNC-CA157 at low μM concentrations, similar to that of alkyl-TPP alone^31, 32^, further modifications will be needed before exploring the utility of this compound in higher organisms (Extended Data Fig. 4b).

### 2-PP modulates the COQ8 active site and KxGQ domain

Targeted inhibition of COQ8 has utility for understanding the CoQ biosynthesis pathway and may present select therapeutic opportunities. However, direct activation of the pathway likely has more immediate benefits. Recently, we discovered that small molecules resembling the CoQ headgroup, including 2-propylphenol (2-PP), enhance COQ8’s ATPase activity through unclear means^21^. A better understanding of how these compounds affect COQ8 could pave the way for more effective CoQ pathway enhancers.

We employed an NMR-based strategy to begin understanding how the CoQ headgroup precursor mimetics stimulate COQ8A ATPase activity. We first validated that COQ8A binds 2-PP by observing reduction of the 2-PP ^1^H NMR signal intensity upon addition of COQ8A^NΔ250^ (Fig. 5a). Next, based on the size of COQ8A^NΔ250^, we opted for a methyl labeling approach amenable to larger proteins to map the protein-ligand interaction. In this scheme, the Ile *δ*1, Leu *δ*1/2, and Val *γ*1/2 methyl groups (collectively ILV) were selectively ^13^C and ^1^H labeled in a perdeuterated background. COQ8A^NΔ250^ contains 84 ILV residues, providing a significant number of probes to detect 2-PP binding. The ^1^H-^13^C HMQC spectrum of apo COQ8A^NΔ250^ showed 85% of the expected peaks, with some not observed due to resonance overlap or broadening (Extended Data Fig. 5b). Through a combination of NOESY and mutational data (Fig. 5c), we assigned at least one methyl group for 39 residues in COQ8A^NΔ250^. Assigned residues formed four distinct clusters in the N-lobe, nucleotide binding site, KxGQ domain, and part of the C-lobe (Fig. 5b, d, Extended Data Fig. 5c). Assignment in the majority of the C-lobe was limited likely because the μs-ms timescale protein dynamics in this region that result in peak broadening. The addition of 1 mM 2-PP led to widespread chemical shift perturbations (CSPs) in the COQ8A^NΔ250^ spectrum, with 33% of the peaks assigned in both apo and 2-PP-bound states passing our significance threshold (Fig. 5e). Significant shifts occurred in all clusters except for the N-lobe and could suggest that 2-PP is exerting an allosteric effect on the protein structure (Fig. 5f). We compared the four assigned clusters by averaging CSP values for residues in these regions (Extended Data Fig. 5c). The N-lobe cluster was essentially unaffected by 2-PP binding, and the most robust changes were observed in the nucleotide binding site and KxGQ domain.

**Fig. 5.**
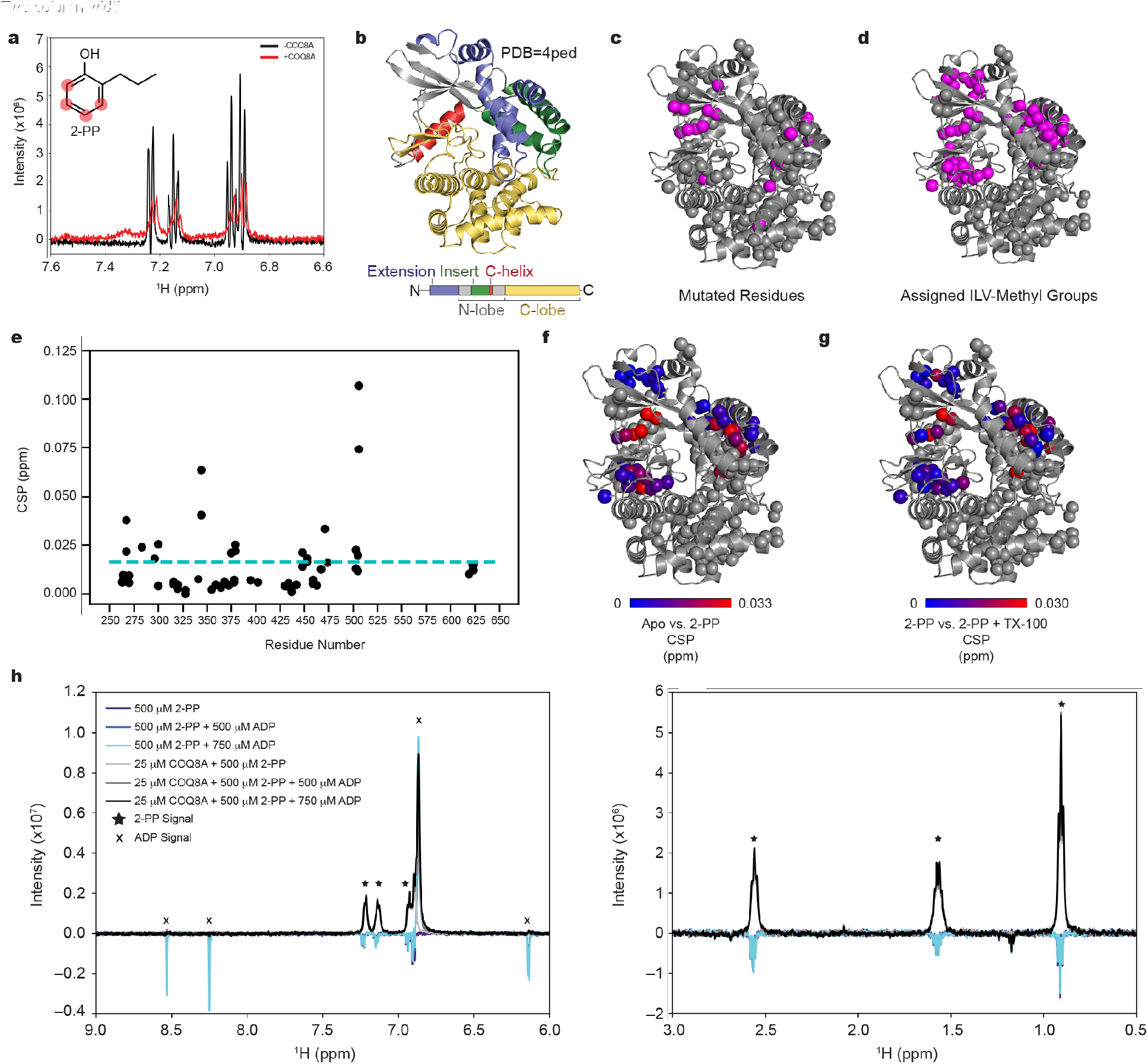
NMR reveals 2-PP induced COQ8A^NΔ250^ structural changes. **a**, ^1^H NMR of the aromatic portion of 2-PP with and without COQ8A^NΔ250^. **b**, Structural overview of COQ8A (PDB=4ped**). c**, Mutated ILV residues for methyl assignment mapped onto 4ped**. d**, Assigned ILV methyls mapped onto 4ped. **e**, 2-PP induced chemical shift perturbations (CSPs) mapped onto the primary COQ8A sequence. The significance threshold (cyan) represents 1.5x the 10% trimmed mean. **f**, 2-PP induced CSPs mapped onto the 4ped structure with the color scale ranging from 0-3σ over the 10% trimmed mean. **e**, Further Triton X-100 induced CSPs mapped onto the 4ped structure with the color scale ranging from 0-3σ over the 10% trimmed mean. **g**, WaterLOGSY spectra of 2-PP and ADP with and without COQ8A^NΔ250^. Spectra are phased such that signals from non-binding compounds are negative.

The large CSPs observed in the nucleotide binding site raised the possibility that 2-PP could also bind in this same location. To address this, we performed WaterLOGSY NMR, a sensitive method for observing ligand-protein interaction^33^. Spectra were collected for a mixture of COQ8A^NΔ250^ and 2-PP before and after addition of ADP. The spectra were phased such that signals from non-binding compounds are negative, while signals arising from binding are positive. Both ADP and 2-PP gave negative WaterLOGSY signals in the absence of protein (purple and blue spectra), and the addition of protein led to either no signal (ADP) or a positive signal (2-PP), both of which are indicative of binding (Fig. 5h). Addition of ADP had no effect on the intensities of the 2-PP WaterLOGSY signals (black and gray spectra), indicating that binding is not competitive. This result demonstrates that 2-PP does not bind in the nucleotide binding site and that observed CSPs in this region are caused by an allosteric effect. Although no precise location of the 2-PP binding site was identified from our NMR experiments, the data suggest the interface of the KxGQ domain and C-lobe as a likely candidate for the binding site since large CSPs were observed. The NMR data also reveal allosteric structural changes induced by 2-PP that extend to the nucleotide binding site.

Stimulated COQ8A^NΔ250^ ATPase activity also requires Triton X-100 (TX-100). We confirmed that TX-100 binds COQ8A^NΔ250^ by using ^1^H NMR (Extended Data Fig. 5a). TX-100 (1 mM) was added to the 2-PP-bound COQ8A sample and induced additional CSPs largely localized to the KxGQ domain (Fig. 5g, Extended Data Fig. 5b). Based on past molecular dynamics simulations of COQ8A-membrane interactions, we suggest that TX-100 could be acting as a membrane mimic to stimulate COQ8A^NΔ250^ ATPase activity. The addition of TX-100 at high protein concentrations led to precipitation of COQ8A^NΔ250^, preventing further investigations beyond this experiment.

### HDX-MS nominates a 2-PP binding site

To further investigate the interaction between COQ8A and 2-PP, we performed hydrogen deuterium exchange mass spectrometry (HDX-MS). HDX-MS is a powerful tool for evaluating solvent accessibility across regions of a protein and can be used to identify putative ligand binding sites and protein conformational changes based on how this accessibility changes in the presence of a ligand^34^. We performed HDX-MS with and without 2-PP. A total of 92 peptides were identified as statistically different. Regions 251-281, 289-306, 366-381, 397-406 showed strong deprotection upon 2-PP binding and regions 287-288, 331-340, 347-365, 382-396, 407-414, 448-468, 482-494, 529-544, 571-581, 607-618, 629-639 showed intermediate deprotection (Fig. 6c). The observed broad-scale deprotection of the protein upon 2-PP binding is consistent with 2-PP inducing an allosteric effect (Fig. 6a, b). Such broad scale deprotection, consistent with increased dynamics or loss of structure of a protein is unusual in our experience. The strongest deprotection was seen in the KxGQ domain, in accord with the significant CSPs observed in this region via NMR.

**Fig. 6.**
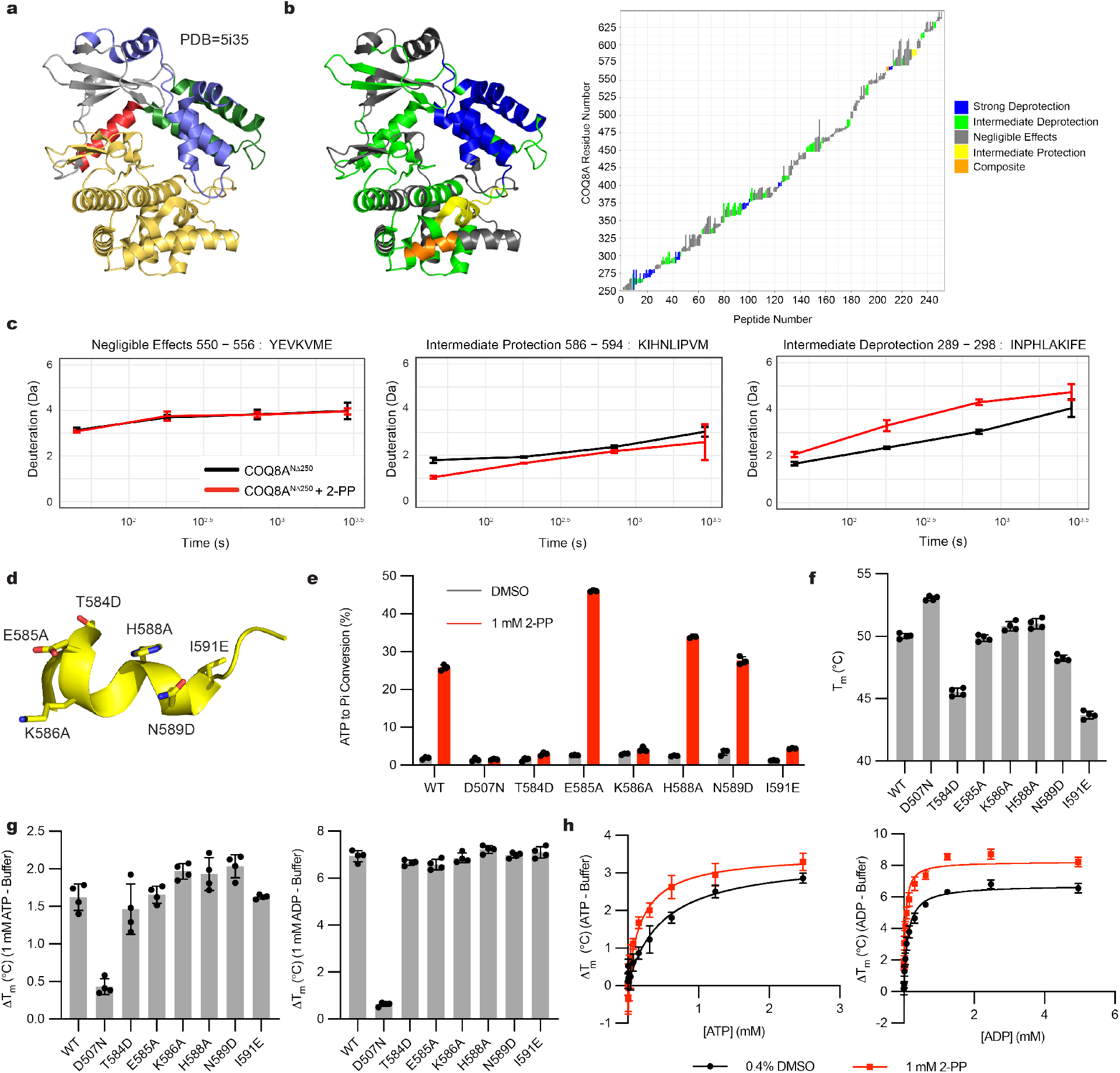
HDX-MS reveals largescale deprotection and candidate 2-PP binding site. **a**, Crystal structure of COQ8A^NΔ254^ (PDB=5i35). **b**, HDX-MS results mapped onto the 5i35 structure and annotated peptides mapped onto the primary COQ8A sequence. **c**, Representative uptake plots for different peptide classifications showing the effect of 2-PP binding to COQ8A^NΔ250^ (n = 3 ± SD). **d**, Region protected upon binding as identified by HDX-MS, highlighting introduced mutations. **e**, ATPase activity of WT or protected region mutants with either DMSO or 1 mM 2-PP (n = 3 ± SD). **f**, Thermal stability of WT COQ8A^NΔ250^ and protected region mutants, as assessed by DSF (n = 4 ± SD). **g**, Thermal stabilization of WT COQ8A^NΔ250^ and protected region mutants by 1 mM ATP or 1 mM ADP, as assessed by DSF (n = 4 ± SD). **h**, Nucleotide binding curves for COQ8A^NΔ250^ with either 1 mM 2-PP or DMSO, as assessed by DSF (n = 4 ± SD).

In addition to this general deprotection, we also found a 13 amino acid region (region 582-594) whose solvent accessibility was reduced upon 2-PP binding (Fig. 6b, c, d). These data suggest that a 2-PP binding pocket is in this region, which is near a hydrophobic patch previously shown to open upon nucleotide binding. Finally, two peptides located in the region 563-570 showed a composite behavior. Importantly, there are several false negatives (peptides showing negligible differences located within the regions mentioned above). This false categorization is most probably due to the strict k-means clustering classification or due to high standard deviation in the HDX measurements.

To test whether the protected region is involved in 2-PP binding, we measured ATPase activity ± 2-PP for WT COQ8A^NΔ250^, the catalytically inactive D507N mutant, and six individual point mutants (Fig. 6e). Each construct exhibited comparable purity and enhanced stability upon nucleotide binding (Fig. 6g), although select mutants had slightly decreased basal melting temperatures (Fig. 6f). Interestingly, mutations to this region had variable effects, with three mutants exhibiting no stimulation upon 2-PP treatment, and another exhibiting a nearly two-fold increase in ATPase activity compared to the WT. Collectively, these results are consistent with this region harboring important residues for 2-PP binding and activation, and suggest that stronger activation may be possible through further ligand modification; however, a deeper structure-function analysis will be required to validate these findings and further elucidate the 2-PP mode-of-action.

Last, we determined the *K_d,app_* for ATP and ADP in the presence of 2-PP by using differential scanning fluorimetry (DSF) (Fig. 6h). We found modest but significant increases in nucleotide affinity with the addition of 2-PP, which could contribute to the observed increase in ATPase activity. Overall, our data indicate that 2-PP stimulates COQ8A ATPase activity by inducing an allosteric modulation of the KxGQ domain and nucleotide binding site. We also implicate the interface between the KxGQ domain and the C-lobe as a potential 2-PP binding site.

## Discussion

In this work, we characterized a series of *in vitro* COQ8A inhibitors, solved an inhibitor-bound crystal structure, specifically targeted our lead candidate to mitochondria, and provided the first inhibition of endogenous COQ8 function in a biological context. Additionally, we determined that the CoQ precursor mimetic 2-propylphenol allosterically modulates the quintessential KxGQ domain and active site of COQ8A to enhance nucleotide binding and ATPase activity. This work sets a foundation for further chemical probe development against COQ8 proteins and the undercharacterized UbiB protein family at large.

Small molecule tools have been instrumental in the biological characterization of PKL family members^2, 35^. Development of specific inhibitors has informed kinase-mediated signaling and resulted in numerous therapeutically relevant molecules^3^. Kinase inhibition often does not phenocopy genetic knockouts, which has also enabled interrogation of protein functions independent of canonical phosphorylation or ATP binding^5, 36^. Given the atypical nature of UbiB proteins and a lack of direct evidence for *trans* phosphorylation, development of chemical tools to modulate their activity in cells is sure to aid in elucidating their precise biological functions.

The level of CoQ in the body has a variety of complex implications in human health^19^. Owing to its extreme hydrophobicity and limited bioavailability^37^, cells are primarily reliant on endogenous production to retain their CoQ pool. Therefore, the ability to manipulate CoQ endogenous biosynthesis presents unique opportunities to affect human health. Nine of the genes required for CoQ production harbor known mutations that result in primary CoQ deficiency, leading to disorders such as ataxias and myopathies^19^. Previously, the overexpression of Coq8p in *S. cerevisiae* has been utilized to overcome deficiencies in several CoQ biosynthetic proteins^38, 39^, although the mechanism remains unclear. Our investigation into 2-PP induced activation sets the foundation for understanding the regulation of COQ8 protein activity at a deeper level, and suggests that the specific activation of COQ8 proteins could have therapeutic benefit against diseases associated with primary and secondary CoQ deficiency.

Although the therapeutic implications for activating CoQ biosynthesis are more obvious than inhibiting this pathway, there are also potential benefits to inhibition^22^. Limitations to CoQ production through genetic or dietary intervention increase lifespan in *C. elegans*^40^ and mice^41^. CoQ is also utilized by the oxidoreductase FSP1 to inhibit iron-mediated cell death in cancer^42, 43^, presenting CoQ limitation as a potential anti-cancer strategy. Furthermore, established disease-related COQ8A mutations are reported to likely be gain-of-function variants^12^, suggesting that COQ8A inhibition could be therapeutically beneficial in select contexts. Last, there is still much to be learned about how the CoQ biosynthetic proteins interact and function, and specific agonistic and inhibitory tools could aid in these investigations. Other inhibitors against CoQ biosynthesis were developed previously, including 4-nitrobenzoate (an inhibitor of COQ2)^44^, clioquinol (an inhibitor of COQ7)^45^, and the osteoporosis drug zoledronic acid^46^. However, previously identified inhibitors are administered on the timescale of days, whereas TPP-UNC-CA157 presents a unique opportunity to perturb acutely and rapidly CoQ biosynthesis. This molecule has immediate use as a research tool to investigate the pathway, but toxicity issues at concentrations that are efficacious will need to be overcome before moving towards clinical application.

In this work, we utilized a triphenylphosphonium moiety to target specifically our small molecule inhibitor to the mitochondrial matrix. This is a well-established strategy for mitochondrial targeting, most prominently for the established mitochondrial antioxidant MitoQ^47^. Given the role of mitochondria as central hubs of metabolism, specific organellar targeting offers a wide range of potential applications to modulate cellular function. However, there is a growing awareness for the toxicity of TPP compounds^31, 32^. Negative impacts of TPP compounds on bioenergetics were reported in the low μM range, including for MitoQ^32^. Efforts are underway to decrease this toxicity by developing modified lipophilic cations^48^, which could inform future efforts to improve our TPP-UNC-CA157 inhibitor.

UbiB proteins are found in all organisms and have established associations with prenyl lipid biology. In eukaryotes, the COQ8 proteins are likely ATPases required for CoQ biosynthesis; however, specifics regarding how this activity is coupled to CoQ production remain unknown^21^. More recently, the yeast intermembrane space UbiB proteins were connected to CoQ distribution throughout the cell, but their precise mechanism of action also remains to be determined^49^. Here we have begun to understand how the activity of these proteins is modified by CoQ precursor mimetics at a structural level. This will inform future models regarding how these proteins interact with CoQ-like molecules and how they function in a cellular context. Additionally, we demonstrate the first inhibition of an endogenous UbiB protein, setting a foundation for future chemical probe and drug development against this understudied protein family.

This integrated approach will help develop small molecule probes to manipulate COQ8 activity *in vitro* and in cells, expanding the molecular toolkit for the PKL superfamily and providing new resources for establishing how auxiliary proteins assist the biosynthesis of CoQ. Beyond COQ8, we anticipate that these chemical tools will form the basis for selecting an even larger set of molecules that will enable mechanistic studies into the functions of the widespread UbiB protein family.

## Supporting information

HDX-MS Supplement

Inhibitor Supplement

NMR Supplement

Synthesis Supplement

## Methods

### COQ8A^NΔ250^ General Expression and Purification

The general purification method has been documented previously^1^. COQ8A^NΔ250^ constructs were overexpressed in *E. coli* by autoinduction^2^. Cells were isolated and frozen at −80 °C until further use. For protein purification, cells were thawed and resuspended in Lysis Buffer [20 mM HEPES (pH 7.2), 300 mM NaCl, 10% glycerol, 5 mM 2-mercaptoethanol (BME), 0.25 mM phenylmethylsulfonyl fluoride (PMSF)] (4 °C). The cells were lysed by sonication (4 °C, 75% amplitude, 20 s x 2). The lysate was clarified by centrifugation (15,000 *g*, 30 min, 4 °C). The cleared lysate was mixed with cobalt IMAC resin (Talon resin) and incubated (4 °C, 1 h). The resin was pelleted by centrifugation (700 *g*, 2 min, 4 °C) and washed four times with Wash Buffer [20 mM HEPES (pH 7.2), 300 mM NaCl, 10% glycerol, 5 mM BME, 0.25 mM PMSF, 10 mM imidazole]. His-tagged protein was eluted with Elution Buffer [20 mM HEPES (pH 7.2), 300 mM NaCl, 10% glycerol, 5 mM BME, 100 mM imidazole]. The eluted protein was concentrated with a MW-cutoff spin filter (50 kDa MWCO) and exchanged into Storage Buffer [20 mM HEPES (pH 7.2), 300 mM NaCl, 10% glycerol, 5 mM BME]. The concentration of 8His-MBP-[TEV]-COQ8A^NΔ250^ was determined by its absorbance at 280 nm (*ε* = 96,720 M^-1^cm^-1^)(MW=89.9 kDa). The fusion protein was incubated with *Δ*238TEV protease (1:50, TEV:fusion protein, mass:mass)(1 h, RT). The TEV protease reaction mixture was mixed with cobalt IMAC resin (Talon resin) and incubated (4 °C, 1 h). The resin was pelleted by centrifugation (700 *g*, 2 min, 4 °C). The unbound COQ8A was collected and concentrated with a MW-cutoff spin filter (30 kDa MWCO) and exchanged into storage buffer. The concentration of COQ8A^NΔ250^ was determined by Bio-Rad Protein Assay according to manufacturer protocol. The protein was aliquoted, frozen in N_2 (l)_, and stored at −80 °C. Fractions from the protein preparation were analyzed by SDS-PAGE. Coq8p^NΔ41^ and mutants were also purified using this general method.

### General 4-anilinoquinoline Synthesis

4-chloroquinoline derivative (150 mg, 0.67 mmol) and aniline (0.74 mmol) were suspended in ethanol (10 mL) and heated to reflux for 16 h. The crude mixture was purified by flash chromatography using EtOAc/hexane followed by 1–5% methanol in EtOAc (or by re-crystallization). After solvent removal under reduced pressure, the product was obtained. All compounds were >98% pure by ^1^H/^13^C NMR and LC-MS and melting points were consistent with previous reports. All compounds in the initial DSF screen have been previously described^3–8^. Full structures for all compounds in the DSF screen can be found in the supplemental information.

### Differential Scanning Fluorimetry

The general DSF method has been documented previously^9^. For screening 4-anilinoquin(az)olines 20 μL reactions containing 10 μM COQ8A^NΔ250^, 100 μM inhibitor from 10 mM stock in DMSO, 5 mM MgCl_2_, 50 mM HEPES pH 7.5, 75 mM NaCl, and 4x Sypro Orange dye (Thermo S6651) were made in MicroAmp Optical 96-well reaction plates (Thermo N8010560), centrifuged (200 *g*, RT, 30 sec) and incubated at room temperature in the dark for 10 min. Fluorescence was then monitored with the ROX filter using a QuantStudio 6 Real-Time PCR system (QuantStuido Real-Time pCR v1.2 software) along a temperature gradient from 4-95 °C at a rate of 0.025 °C/s. Protein Thermal Shift software v1.3 (Applied Biosystems) was used to determine T_m_ values by fitting to a Boltzmann model. Melt curves flagged by the software were manually inspected. Two data points were omitted after inspection (UNC-CA331, UNC-CA310) resulting in only duplicate measurements reported for those compounds. Each inhibitor was run in triplicate and error bars represent SD. *D*T_m_ values were determined by subtracting the average of matched vehicle controls from the same plate. UNC-CA157 binding to COQ8A^NΔ250^ and Coq8p^NΔ41^ WT and active site mutants were also analyzed using this method.

To determine the inhibitor *K_d,app_*, 20 μL reactions contained 1 μM COQ8A^NΔ250^, 0-200 μM inhibitor with a final concentration of 2% DMSO, 4 mM MgCl_2_, 100 mM HEPES pH 7.5, 150 mM NaCl, and 4x Sypro Orange dye. Samples were run and analyzed as described above. *D*T_m_ values were plotted against [inhibitor] in GraphPad Prism Version 8.4.3 and fit to nonlinear regression (one site specific binding) to determine *K_d,app_* values. The experiments were performed in triplicate and error bars represent SD. Error values in *K_d,app_* values represent 95% confidence intervals.

To determine the nucleotide *K_d,app_*, 20 μL reactions contained 1 μM COQ8A^NΔ250^, either 0-4.96 mM ADP (Sigma A2754) or 0-2.48 mM ATP (Sigma A2383), and either 1 mM 2-PP or 0.4% matched DMSO. The other reaction components were 4 mM MgCl_2_, 100 mM HEPES pH 7.5, 150 mM NaCl, and 5x Sypro Orange dye. Samples were run and analyzed as described above. *K_d,app_* values were determined as described for inhibitors. Experiments were performed in quadruplicate and error bars represent SD. Error values in *K_d,app_* represent 95% confidence intervals. Nucleotide binding experiments for COQ8A^NΔ250^ WT and mutants at 1 mM ATP and 1 mM ADP were also performed using this method with no DMSO vehicle (n=4, error bars represent SD).

### ATPase Assay

ATPase assays for COQ8A^NΔ250^ were performed essentially as described in Reidenbach et al.^10^ The reaction was initiated by adding a mixture of ATP (Promega V703), 2-propylphenol (Sigma W352209), Triton X-100 (Sigma T9284), and indicated inhibitors from stocks in DMSO (Sigma D2650) in Reaction Buffer [100 mM HEPES pH 7.5, 4 mM MgCl_2_, 150 mM NaCl] to purified COQ8A^NΔ250^ also in reaction buffer. Final reaction conditions were 100 μM ATP, 1 mM 2-PP, 0.5 μM COQ8A^NΔ250^, 1 mM Triton X-100, and 0-50 μM indicated inhibitor with equivalent DMSO vehicle. The final reaction volume was 15 μL. Reactions were performed in clear 384 well plates (Fisher 12565506). Once the reactions were started and mixed, plates were sealed and incubated at 30°C for 45 min. Reactions were then quenched with 35 μL of Cytophos^TM^ reagent (Cytoskeleton, Inc. BK054), incubated at RT for 10 min, and absorbance was read at 650 nm in a Biotek Cytation 3 plate reader. Absorbance was converted to concentration of inorganic phosphate with a 0-50 μM standard curve run in parallel using the phosphate standard provided in the Cytophos^TM^ kit diluted in reaction buffer. Experiments were performed in triplicate and error bars on the graph represent SD. To determine IC_50_ values the data were plotted in GraphPad Prism Version 8.4.3 and fit to a nonlinear regression ([inhibitor] vs. response, variable slope, four parameters). Indicated error in IC_50_ values represents 95% confidence intervals. For comparisons between COQ8A mutants in the HDX-MS protected region, the same ATPase assay was performed with final reaction conditions of 100 μM ATP, 0.5 μM COQ8A^NΔ250^, 1 mM Triton X-100, and either 1 mM 2-PP or matched DMSO in Reaction Buffer.

### Cell Transfections and BRET Measurements

For COQ8A and COQ8B cellular BRET measurements, N-or C-terminal NanoLuc/Kinase fusions were encoded in pFN31K expression vectors (Promega), including flexible Gly-Ser-Ser-Gly linkers between Nluc and each full-length kinase. The COQ8A and COQ8B open reading frames both corresponded to UNIPROT Isoform 1 (Q8NI60 and Q96D53, respectively). The N-terminal fusions to NanoLuc to COQ8A and COQ8B were found to be optimal for BRET and used for assay development and compound potency testing. HEK-293 cells were transfected with NanoLuc-COQ8A or NanoLuc-COQ8B fusion constructs using FuGENE HD (Promega) according to the manufacturer’s protocol. Briefly, fusion constructs were diluted together into Transfection Carrier DNA (Promega) at a mass ratio of 1:9 (mass/mass/mass) in Opti-MEM (Gibco), after which FuGENE HD was added at a ratio of 1:3 (μg DNA: µL FuGENE HD). For example, for a 1mL size transfection complex, 1µg of the NanoLuc fusion DNAs was combined with 9µg Transfection Carrier DNA in 1 mL of Opti-MEM. 1 part (vol) of FuGENE HD complexes thus formed were combined with 20 parts (vol) of HEK-293 cells suspended at a density of 2 x 10^5^ per mL in Opti-MEM containing 1% (v/v) FBS. Transfected cell suspension were seeded at 100 µL per well (20,000 cells/well) into white tissue culture-treated assay plates (Corning CAT# 3917) followed by incubation in a humidified, 37°C/5% CO_2_ incubator for 18−24 hr. The total concentrations for NanoLuc® fusion plasmids were 5 ng/well, and the total concentration of DNA was 50 ng/well. All chemical inhibitors were prepared as concentrated stock solutions in DMSO (Sigma-Aldrich) and diluted in Opti-MEM (unless otherwise noted) to prepare working stocks. Cells were equilibrated with the COQ8 BRET probe and test compound prior to BRET measurements, with an equilibration time of 2 hours unless otherwise noted. COQ8 BRET probe was prepared first at a stock concentration of 100X in DMSO, after which the 100X stock was diluted to a working concentration of 20X in BRET probe dilution buffer (12.5 mM HEPES, 31.25% PEG-400, pH 7.5). For COQ8 BRET probe dose response measurements, the COQ8 BRET probe was added to the cells in an 8 point, 2-fold dilution series starting at a final concentration of 1µM. For target engagement analysis, the COQ8 BRET probe was added to the cells at a final concentration of 1µM for COQ8A and 0.25µM for COQ8B. To measure BRET with the COQ8 BRET probe, NanoBRET Target Engagement Substrate (Promega) and Extracellular NanoLuc Inhibitor (Promega) were added according to the manufacturer’s recommended protocol, and filtered luminescence was measured on a GloMax Discover luminometer equipped with 450 nm BP filter (donor) and 600 nm LP filter (acceptor), using 0.5 s integration time. Raw BRET ratios were calculated by dividing the acceptor counts by the donor counts. Milli-BRET units (mBU) were calculated by multiplying the raw BRET values by 1000.

Apparent BRET probe affinity values (EC_50_) were determined using the sigmoidal dose-response (variable slope) equation available in GraphPad Prism (Equation 1);

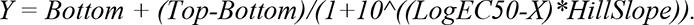

For determination of test compound potency, competitive displacement data were plotted with GraphPad Prism software and data were fit to Equation 1 to determine the IC_50_ value.

### HAP1 Cell Growth and Lipid Extraction

WT, COQ8A^KO^, COQ8B^KO^, and COQ8A/B^DKO^ HAP1 cells were obtained from Horizon Discovery and previously validated (Rensvold, 2022). Three 10 cm plates were seeded with 1.4×10^6^ cells for each cell line in IMDM (Thermo 12440053) supplemented with 10% heat inactivated FBS (HI FBS)(Atlanta Biologicals) and 100 U/mL Penicillin-Streptomycin (Pen/Strep)(Thermo 15140122). Cells were grown at 37 °C, 5% CO_2_ for three days to approximately 90% confluency. Each plate of cells was washed with 5 mL dPBS then isolated by trypsinization. Cells were pelleted (173 *g*, 5 min, RT) and media was removed. Cells were then resuspended in 5 mL media and counted using the Muse Count and Viability Assay (Luminex). 1.2×10^7^ viable cells were collected per sample and pelleted by centrifugation (173 *g*, 5 min, RT). Media was removed and cells were resuspended in 5 mL dPBS. Cells were pelleted again by centrifugation and dPBS was removed. Cell pellets were resuspended in 600 μL cold methanol with 0.1 μM CoQ_8_ (Avanti Lipids 900151p-1mg) as the internal standard. Suspended cells were transferred to screw cap tubes and vortexed for 10 min at 4 °C to lyse. 400 μL cold petroleum ether was added to each tube. Samples were vortexed for 3 min at 4 °C then centrifuged at 800 *g*, 3 min, RT to separate the layers. The top petroleum ether layer was collected in a new tube and the extraction was repeated with an additional 400 μL petroleum ether. The two extractions were combined, and the ether was removed under argon gas. Once dried, the lipids were resuspended in 100 μL methanol, transferred to glass autosampler vials, and submitted for LC-MS lipid measurements.

### Seahorse Mito Stress Test

WT, COQ8A^KO^, COQ8B^KO^, and COQ8A/B^DKO^ HAP1 cells were split, counted via Muse Count and Viability Assay, and 40,000 viable cells per well were plated to a poly-D-lysine-coated XF96 Seahorse plate. Cells were allowed to adhere to the plate overnight at 37 °C, 5% CO_2_ in IMDM supplemented with 10% heat inactivated FBS and 100 U/mL penicillin-streptomycin. After 24 h, the medium was aspirated, cells were washed once with dPBS, and medium was replaced with 180 μL Seahorse XF Base Medium (Agilent 103252-100), pH 7.4, supplemented with 25 mM glucose and 2 mM glutamine. The plate was then incubated for 1 h at 37 °C in a non-CO_2_ incubator. Oxygen-consumption rates (OCR) and extracellular acidification rate were then monitored on a Seahorse XF96 Extracellular Flux Analyzer basally, and in the presence of a Seahorse XF Mito Stress Test (Agilent 10315-100). For the stress test, cells were treated with oligomycin (2 μM final concentration), FCCP (0.5 μM final concentration), and rotenone and antimycin A (0.5 μM final concentration). After the assay, cells were fixed with 1% glutaraldehyde and stained with 1.5% crystal violet, and, after release of the stain with 10% acetic acid, each well was read at an absorbance of 590 nm^11^. OCR measurements for each well were normalized to relative crystal violet absorbance. OCR measurements were plotted and analyzed using GraphPad Prism Version 8.4.3 (n=11-12, error bars represent SD).

### COQ8A^NΔ254^ Crystallography

COQ8A^NΔ254^ was expressed and purified as described for COQ8A^NΔ250^. After the final buffer exchange the protein was concentrated to less than 1 mL. The sample was centrifuged (20,000 *g*, 10 min, 4°C) and the soluble portion was filtered through a 0.22 μm filter. The protein was further purified using a HiLoad 16/600 Superdex 75 pg gel filtration column (GE) with an isocratic elution at 1 mL/min. The protein was exchanged into Crystallography Buffer [50 mM NaCl, 5 mM HEPES pH 7.5, 0.3 mM TCEP] during the FPLC run. Fractions were analyzed by SDS-PAGE. Pure fractions were pooled, concentrated with a MW-cutoff spin filter (30 kDa MWCO), aliquoted, frozen in N_2 (l)_, and stored at −80°C for crystallization trials.

Crystallization screening and optimization was carried out in MRC SD2 plates using a TTP Labtech Mosquito using 50 microliters of reservoir solution. Mixtures of 0.39 mM COQ8A^NΔ254^ and 0.5 mM CA157 was incubated for 30 minutes at ambient temperature before crystallization setup.

Crystals were screened for optimum diffraction at sectors 21 and 23 of the Advanced Photon Source. Data sets for refinement were collected at GM/CA@APS on 2019-12-12 using JBluIce-EPICS^12^ beamline control software. Data was reduced using autoPROC^13^ and XDS^14^. The structures were solved by molecular replacement using PDB 5i35 in Phaser^15^. Geometric restraints for CA157 were prepared with phenix.elbow^16^. Structures were iteratively improved using model building in Coot^17^ refinement in Phenix^18^, and sterochemical validation in MOLPROBITY^19^. Final models were prepared for deposition using PDB_EXTRACT^20^. SBGrid provided curated crystallographic software^21^.

#### 7UDP

Optimal crystals were obtained by mixing 200 nL COA8A-CA157 with 250 nL of 0.65 M sodium succinate and 0.1 M HEPES buffer, pH 8. Data was collected at 100K on APS beamline 23ID-D with 1.0332 Å X-rays, on a Dectris Pilatus3-6M detector at 350 mm sample-detector distance, 1800 frames, 0.2 degree per frame.

#### 7UDQ

Optimal crystals were obtained by mixing 200 nL COA8A-CA157 with 250 nL of 0.65 M sodium succinate, pH 7. Data was collected at 100K on APS beamline 23ID-B with 1.0331 Å X-rays, using an Eiger 16M at 240 mm sample-detector distance, 1800 frames, 0.2 degree per frame.

### Site Directed Mutagenesis

The COQ8A^NΔ250^, COQ8A^NΔ254^, and Coq8p^NΔ41^ in pVP68K vectors have been described previously^1,2^^2^. Point mutations were introduced via PCR-based mutagenesis. Reactions were DpnI digested and transformed into DH5*α* competent *E. coli* cells. Plasmids were isolated from transformants and Sanger sequencing was used to identify those harboring the correct mutations. Oligonucleotides were purchased from IDT (Coralville, IA, USA).

### TPP-UNC-CA157 Synthesis

TPP-UNC-CA157 was synthesized in collaboration with the Washington University Center for Drug Discovery. Detailed synthesis methods can be found in the Supplemental Information. This general synthetic strategy for mitochondrial targeting has been described previously^23^.

### *De novo* ^13^C-CoQ_10_ Production

This general strategy for tracking *de novo* CoQ production was used for yeast in Reidenbach et al^10^ and was optimized here for HAP1 cells. WT, COQ8A^KO^, COQ8B^KO^, and COQ8A/B^DKO^ HAP1 cells were split, counted via Muse Count and Viability Assay, seeded in 10 cm plates at 1.5×10^6^ cells per plate in IMDM with HI FBS and Pen/Strep, and allowed to grow at 37°C, 5% CO_2_ for 48 h. Cells were then rinsed with 5 mL dPBS, swapped into 10 mL DMEM (no glucose) (Thermo 11966025) supplemented with HI FBS, Pen/Strep, and 10 mM galactose, and grown again at 37°C, 5% CO_2_. After 24 h cells were swapped into the same media containing 10 μM 4-hydroxybenzoate-(*phenyl*-^13^C_6_) (Sigma 587869) and either indicated inhibitor or matched DMSO vehicle. Cells were incubated for 1 h at 37°C, 5% CO_2_. Cells were washed with dPBS then isolated by trypsinization. Cells were counted via Muse Count and Viability Assay and equivalent viable cells were collected in each experiment (UNC-CA157=2.92×10^6^ cells; TPP-UNC-CA157=7.91×10^6^ cells; TPP-UNC-CA157 dose response=4.53×10^6^ cells) via centrifugation (300 *g*, 5 min). Cells were washed with dPBS then again isolated by centrifugation (300 *g*, 5 min) and dPBS was removed. Cell pellets were frozen in N_2 (l)_ and stored at −80 °C. On the day of analysis cell pellets were thawed on ice. Lipids were extracted and dried under Ar_(g)_ as described above for untreated HAP1 cells. Dried lipids were resuspended in an equivalent volume of methanol per experiment (UNC-CA157=60 μL; TPP-UNC-CA157=100 μL; TPP-UNC-CA157 dose response=60 μL), then submitted for LC-MS lipid measurements.

### LC-MS Lipid Measurements

A Vanquish Horizon UHPLC system (Thermo Scientific) coupled to an Exploris 240 Orbitrap mass spectrometer (Thermo Scientific) was used for LC-MS analysis. A Waters Acquity CSH C18 column (100 mm × 2.1 mm, 1.7 µm) was held at 35°C with the flow rate of 300 µL/min for the separation of the extracted lipids. A Vanquish binary pump system was employed, and mobile phase A consisted of 5 mM ammonium acetate in ACN/H_2_O (70/30, v/v) containing 125 µL/L acetic acid, while mobile phase B consisted of 5 mM ammonium acetate in IPA/ACN (90/10, v/v) with the same additive. The gradient was set as follows: B was held at 2% for 2 min and ramped up to 30% over the next 3 min, before further increased to 50% within 1 min and to 85% over 14 min, and then raised to 99% over 1 min and held for 4 min. The column was re-equilibrated for 5 min at 2% B before the injection of next sample. Samples were ionized by a heated ESI source with a vaporizer temperature of 350°C. Sheath gas was set to 50 units, auxiliary gas was set to 8 units, sweep gas was set to 1 unit. The ion transfer tube temperature was kept at 325°C with 70% RF lens. Spray voltage was set to 3,500 V for positive mode and 2,500 V for negative mode. Each sample was injected twice to be analyzed in positive and negative modes separately. Full MS scans were acquired at 120,000 resolution from m/z 200 to 1,700 with in-source Easy-IC enabled. Normalized AGC target was set to 100% with the maximum ion injection time of 50 ms. Targeted quantitative analysis of CoQ_10_, PPHB_10_, ^13^C_6_-CoQ_10_, ^13^C_6_-PPHB_10_ and internal standard CoQ_8_ was processed using TraceFinder 5.1 (Thermo Scientific) with the mass accuracy of 5 ppm. The result of peak integration was manually examined. Peak areas were normalized to the CoQ_8_ internal standard. Experiments were performed in triplicate and error bars represent SD.

### HAP1 Growth Assay

WT HAP1 cells were split and counted via Muse Count and Viability Assay. 2×10^4^ cells were seeded per well in a 96 well plate (Greiner Bio-One 655090). Cells were allowed to adhere for 1 h at RT in the hood, then incubated at 37°C, 5% CO_2_ overnight. The next day cells were washed once with dPBS then swapped into DMEM (no glucose) (Thermo 11966025) supplemented with HI FBS, Pen/Strep, either 10 mM glucose or galactose, and TPP-UNC-CA157 at indicated concentrations with equivalent DMSO vehicle. Cells were incubated at 37°C, 5% CO_2_ for 90 h. Cell growth was monitored using a Sartorius IncuCyte S3 Live-Cell Analysis System at 10x magnification in the phase channel. Percent confluency and associated error were determined using the Incucyte S3 Software. Graphs were made using GraphPad Prism Version 8.4.3 (n=6, error represents SEM).

### 1D Ligand-Observed NMR

COQ8A^NΔ250^ was purified with the general method described above. After the final buffer exchange the protein was concentrated to less than 1 mL. The sample was centrifuged (20,000 *g*, 10 min, 4 °C) and the soluble portion was filtered through a 0.22 μm filter. The protein was further purified using a HiLoad 16/600 Superdex 75 pg gel filtration column (GE) with an isocratic elution at 1 mL/min. The protein was exchanged into Analytical Buffer [100 mM NaCl, 20 mM HEPES, pH 7.2] during the FPLC run. Fractions were analyzed by SDS-PAGE and concentrated for NMR analysis. Binding of 2-PP and TX-100 was assessed by ^1^H NMR. Samples consisted of 200 μM ligand, 20 μM DSS, 8% D_2_O, 100 mM NaCl, 20 mM HEPES, pH 7.2. Spectra were collected at 30 °C on a Bruker Avance III spectrometer running Topspin version 3.5pl7 and operating at 500 MHz (^1^H). After collecting spectra without protein, a final concentration of 22.8 μM COQ8A^NΔ250^ was added to the sample and spectra were collected again.

For WaterLOGSY experiments COQ8A^NΔ250^ was expressed and purified using the general method and stored at −80 °C. Protein was thawed on ice and exchanged four times into dPBS (Thermo 14190250) using a 30 kDa MW-cutoff spin filter to remove HEPES and glycerol. It was then further dialyzed for 2 h, then again overnight using a Slide-A-Lyzer Mini Dialysis Device (20 kDa MWCO, 0.5 mL) (Thermo 88402) based on manufacturer protocol. The sample was stored at 4 °C until NMR analysis. Samples were prepared by making a 2X stock solution containing 1 mM 2-PP (from stock in DMSO-d6), 10% D_2_O. Half of the stock solution was diluted to a final concentration of 500 μM 2-PP, 5% D_2_O. An aliquot of COQ8A^NΔ250^ stock was added to the other half, then brought to a final concentration of 25 μM COQ8A, 500 μM 2-PP, 5% D_2_O. ADP was titrated into final concentrations of 500 and 750 μM using a 100 mM stock. DSS was added to each sample at the end of each titration for referencing. The final samples contained 0.08% DMSO-d6. WaterLOGSY experiments were recorded at 10 °C on a Bruker Avance III spectrometer operating at 600 MHz (^1^H) using Topspin 3.5pl7. Experiments were recorded using the Bruker pulse program ephogsygpno.2 with a mixing time of 1.5 s and 2 s recycle delay. A 40 ms T_1_*ρ* filter was used to suppress the protein signals. All 1D spectra were processed with 1 Hz exponential line broadening using nmrPipe^24^ and visualized using the nmrglue python package^25^.

### 13C-Labeled COQ8A^NΔ250^ Expression and Purification

The ^13^C labeling schemes have been described previously^26–28^. *E. coli* harboring the COQ8A^NΔ250^ PVP68K plasmid were used to inoculate 20 mL LB cultures with 50 mg/L kanamycin (KAN) and 15 mg/L chloramphenicol (CAM). Cultures were grown at 37 °C, 220 RPM until OD_600_=0.8-1. Then 5 mL of the LB culture was used to inoculate a 50 mL culture of M9/H_2_O minimal media [12.8 g/L Na_2_HPO_4_•7H_2_O, 3g/L KH_2_PO_4_, 0.5 g/L NaCl, 1 g/L NH_4_Cl, 4 g/L glucose, 2 mM MgSO_4_, 0.1 mM CaCl_2_, 0.5 mg/L FeSO_4_, 5 mg/L thiamine, 0.5X BME vitamin solution (Sigma B6891), KAN, CAM, pH=7.4 in H_2_O]. M9/H_2_O cultures were grown until OD_600_=0.8-1. Cells were pelleted by centrifugation (4000 *g*, 15 min, RT) and spent media was decanted. Cell pellets were then resuspended in 100 mL M9/D_2_O minimal media [6.8 g/L Na_2_HPO_4_, 3 g/L KH_2_PO_4_, 0.5 g/L NaCl, 4 g/L deuterated glucose (Sigma 552003), 0.25 g/L ISOGRO-15N, D Powder (Sigma 608300), 1 g/L ^15^NH_4_Cl (Sigma 299251), 2 mM MgSO_4_, 0.1 mM CaCl_2_, KAN, CAM, pH=7.4, in D_2_O (Sigma 756822)]. 100 mL M9/D_2_O minimal media cultures were grown at 37 °C until OD_600_=0.8-1. The 100 mL cultures were then diluted in 400 mL M9/D_2_O minimal media to yield a final 500 mL culture. 500 mL cultures were grown at 37 °C. At 30 min prior to induction (OD_600_=0.75) labeling precursors were added to the cultures. ^13^C-ILV labeling used 150 mg/L 2-Keto-3-(methyl-d_3_)-butyric acid-4-^13^C,3-d sodium salt (Sigma 691887) and 70 mg/L 2-Ketobutryic acid-4-^13^C,3,3,d_2_ sodium salt hydrate (Sigma 589276) as the precursors. Dimethyl LV labeling for geminal pair determination used 150 mg/L 2-Keto-(3-methyl-^13^C)-butyric-4-^13^C,3-d acid sodium salt (Sigma 589063). Valine only labeling used 150 mg/L 2-Keto-3-(methyl-d_3_)-butyric acid-4-^13^C,3-d sodium salt (Sigma 691887) and 150 mg/L sodium 4-methyl-2-oxovalerate (Sigma K0629). Cultures were grown for another 30 min at 37 °C to reach OD_600_=0.8-1. Cultures were cold shocked at 4 °C for 10 min, then induced with a final concentration of 100 μM IPTG. Protein was expressed at 25 °C for 20 h. Cells were then isolated and frozen at −80 °C until further use. Proteins were generally purified using the above method with the following changes. Protein preparations for NMR used no glycerol. After the final COQ8A concentration, the protein was exchanged into Analytical Buffer [100 mM NaCl, 20 mM HEPES, pH=7.2] using the 30 kDa MW-cutoff spin filter. Protein was quantified by its absorbance at 280 nm (*ε* = 28,880 M^-1^cm^-1^)(MW=45.6 kDa) and stored at 4 °C until NMR analysis.

### Protein-Observed NMR

All 2D-4D spectra of COQ8A were recorded at 30°C in 20 mM HEPES, 100 mM NaCl, pH 7.2. 5-10% D_2_O was added for the lock and 50 μM DSS was added for referencing. Most spectra were recorded on a Bruker Avance III spectrometer operating at 750 MHz (^1^H) and equipped with a cryogenic probe. The 4D HMQC-NOESY-HMQC^29^ was recorded with 8 scans per FID with 36 (^13^C) x 28 (^1^H) x 36 (^13^C) x 1024 (^1^H, direct) complex points non-uniformly sampled at 19%. The spectrum was reconstructed using the SMILE algorithm implemented in nmrPipe^30^. The NOE mixing time was 300 ms. The total experimental time was ∼7 days.

3D NOESY-HMQC were collected on an 800 MHz (^1^H) Varian spectrometer running vnmrj version 4.2 and equipped with a cryogenic probe. Two were recorded on a dimethyl-labeled sample of apo COQ8A where both C*γ*s and C*δ*s are labeled per V/L to establish geminal methyl pairing. These spectra were recorded with 98 (^13^C) x 98 (^13^C) x 1024 (^1^H, direct) complex points and mixing times of 50 and 200 ms. The experiment with a 50 ms mixing time used 16 scans per FID while 32 scans were used for the 200 ms experiment. A third 3D NOESY-HMQC was recorded for COQ8A in the presence of 2 mM AMPPNP. This spectrum was recorded in two blocks, both with 68 (^13^C) x 68 (^13^C) x 1024 (^1^H, direct) complex points and a mixing time of 350 ms. The first block consisted of 32 scans per FID while the second block consisted of 16 scans per FID. All NOESY spectra were processed in nmrPipe using the SMILE algorithm^30^. The two blocks of the NOESY spectrum with AMPPNP were co-added using nmrPipe after NUS reconstruction.

All spectra were analyzed using ccpNmr Analysis version 2.4.2^31^. NOESY peak picking was performed manually. Chemical shift assignment was performed using the NOESY data in a ‘methyl walk’ approach^32^, described in more detail in the Supplementary Information. Chemical shift perturbations (CSPs) were calculated according to the formula:

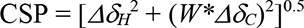

where *Δδ* is the difference in chemical shift for a given residue between apo and 2-PP-bound states and *W* a chemical shift weighting factor (W_C_ = |*γ*_C_/*γ*_H_| = 0.251).

### HDX-MS of COQ8A^NΔ250^ with 2-PP

The protein was expressed and purified by using the above method and stored at −80 °C. At the time of HDX, the protein was thawed on ice and exchanged into Analytical Buffer [100 mM NaCl, 20 mM HEPES, pH = 7.2] by using a 30 kDa MW-cutoff spin filter. Protein concentration was quantified using the Bio-Rad Protein Assay. HDX experiments were performed similarly as described elsewhere^33^. Briefly, automated HDX-MS experiments were carried out using a LEAP Technologies HDX PAL DHR robot (Carrboro, NC) with a three-valve configuration system coupled to a Bruker MaXis Q-ToF operating in positive-ion mode. HDX was initiated by diluting 9 μL of purified COQ8A (15 μM in 20 mM HEPES, 100 mM NaCl, pH 7.2) with 81 μL of 20 mM HEPES, 100 mM NaCl, 5% DMSO, pD 7.2 in D_2_O (Cambridge Isotope Laboratories, Inc., Tewksbury, MA) with and without 8 mM 2-propylphenol. pD was directly measured with a glass electrode and corrected for the deuterium isotope effect (pD = pH + 0.40)^34^. Samples were labeled in triplicate for each labeling time (45, 180, 720 and 2880 s) at 25 °C. Immediately following, 85 μL of each labeled solution was quenched with 85 μL precooled quench buffer (6 M Urea, 10 mM PBS, pH 2.4) at 1 °C. Non deuterated controls were prepared identically but using 20 mM HEPES, 100 mM NaCl, 5% DMSO, pH 7.2. Quenched samples were injected into a temperature-controlled chromatography cabinet maintained at 0 °C throughout the whole experiment to reduce back-exchange. In-line digestion was performed by passing the quenched sample through an custom immobilized pepsin column(2.1 x 50 mm) maintained at 10 °C with 0.1% formic acid in H_2_O for 180 s at a flow rate of 0.2 mL min^-1^. Peptic peptides were trapped and desalted on a ZORBAX 300SB-C8 trap (2.1 × 12.5 mm, 5 µm particles) with 0.1% formic acid in H_2_O for 60 s at a flow rate of 0.2 mL min^-1^. Peptides were separated by using reversed phase chromatography with a XSelect CSH C18 column (2.1 × 50 mm, 2.5μm, Waters, Manchester, UK).

HDX-MS data was analyzed using HDExaminer (version 3.2.1, Sierra Analytics, Modesto, CA). Adjustment of LC retention times and curation of the data were performed manually on all peptides. Non deuterated COQ8A^NΔ250^ peptic peptides were identified using DIA CID MS/MS prior to conducting HDX. Inline digestion of COQ8A^NΔ250^ resulted in 100% sequence coverage. Differential HDX data were evaluated for significant differences by using a hybrid significance test^35^. Normalized HDX differences were clustered^33^ to define thresholds for strong, moderate, and negligible HDX differences. In supplemental data is a volcano plot for identification of statistically different peptides and magnitude of the effects. Negligible (black), moderate (yellow) and strong (blue) effects was revealed by k-mean clustering of the data. Visualization of HDX differences was done plotting clustered significant differences into the COQ8A structure (PDB=5i35)^22^.

## Acknowledgements

We thank members of the Pagliarini lab for helpful discussions throughout this project. This work was supported by NIH awards R35GM131795 (D.J.P.), T32GM008505 (N.H.M.), R24GM136766 (M.L.G.), and R35GM141748 (K.H.W.). This work was also supported by NSF DGE-1747503 (N.H.M.). This study made use of the National Magnetic Resonance Facility at Madison, which is supported by NIH grant R24GM141526. We thank Marco Tonelli and Paulo Falco Cobra fro assistance with NMR data acquisition. We would like to thank Katherine Overmyer, Brett Paulson, Edna Trujillo, and Joshua Coon for assistance with LC-MS method development. This study made use of the Washington University in St. Louis Genome Engineering and iPSC Center for instrument use and the Washington University in St. Louis Center for Drug Discovery for synthesis services.

The SGC is a registered charity (number 1097737) that receives funds from AbbVie, Bayer Pharma AG, Boehringer Ingelheim, Canada Foundation for Innovation, Eshelman Institute for Innovation, Genome Canada, Innovative Medicines Initiative (EU/EFPIA) [ULTRA-DD grant no. 115766], Janssen, Merck KGaA Darmstadt Germany, MSD, Novartis Pharma AG, Ontario Ministry of Economic Development and Innovation, Pfizer, São Paulo Research Foundation-FAPESP, Takeda, and Wellcome [106169/ZZ14/Z]. The US National Institutes of Health (NIH) is acknowledged for support (1U24DK11604-01). We thank Biocenter Finland/DDCB for financial support and the CSC-IT Center for Science Ltd. (Finland) for allocation of computational resources. We also thank Dr. Brandie Ehrmann and Ms. Diane E. Wallace for for LC-MS/HRMS support provided by in the Mass Spectrometry Core Laboratory at the University of North Carolina at Chapel Hill. The core is supported by the National Science Foundation under Grant No. (CHE-1726291).

This research used resources of the Advanced Photon Source, a U.S. Department of Energy (DOE) Office of Science User Facility operated for the DOE Office of Science by Argonne National Laboratory under Contract No. DE-AC02-06CH11357. GM/CA@APS has been funded by the National Cancer Institute (ACB-12002) and the National Institute of General Medical Sciences (AGM-12006, P30GM138396). The Eiger 16M detector at GM/CA-XSD was funded by NIH grant S10 OD012289. With thank Craig Ogata for beamline support. Use of the LS-CAT Sector 21 was supported by the Michigan Economic Development Corporation and the Michigan Technology Tri-Corridor (Grant 085P1000817). The Collaborative Crystallography Core in the Department of Biochemistry, UW-Madison received support from the Department of Biochemistry endowment.

## Data Availability

ILV methyl chemical shifts for COQ8A in the apo state and in the presence of 1 mM 2-PP will be made available on BMRB upon publication. Raw NMR data is being deposited in BMRBig. Raw LC-MS data for lipid measurements are available from the corresponding author on request. Both crystal structures have been deposited in the PDB (7UDP and 7UDQ).

## Author contributions

N.H.M. and D.J.P. wrote the manuscript and conceived the overall project and its design. A.L. and K.H.W. designed the NMR experiments, which were performed and analyzed by A.L. C.R.M.A. performed compound synthesis and contributed to inhibitor development. J.P.R.P. and M.L.G. designed the HDX-MS experiments, which were performed and analyzed by J.P.R.P. Z.F. performed and analyzed mass spectrometry experiments. N.P. contributed reagents (cloning) and developed DSF methods. R.W.S. and C.A.B. performed crystallization trials and solved the crystal structure. J.D.V., C.A.Z., C.R.C., and M.B.R. performed NanoBRET analyses. All authors reviewed and edited the manuscript.

## Supplementary Information

Supplementary Information is available for this paper.

- Structures of all compounds, IC_50_ values, and *K_d,app_* values for Figure 1
- Synthesis information for TPP-UNC-CA157
- NMR methyl assignment process
- HDX-MS analysis figures

**Correspondence and requests for materials** should be addressed to D.J.P.

## Extended data

**Extended Data Fig. 1.**
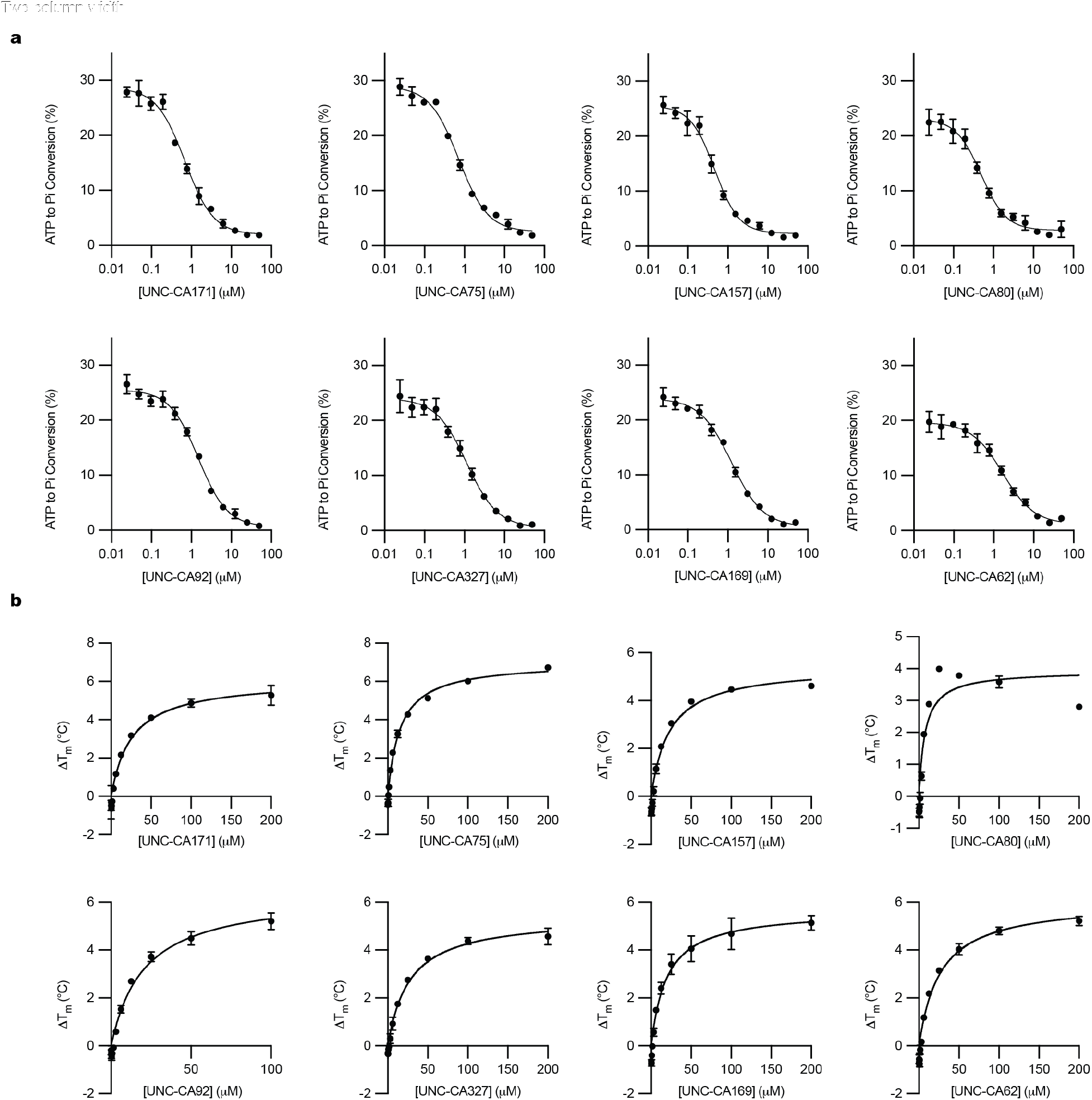
COQ8A inhibition *in vitro*. **a**, Inhibition of COQ8A^NΔ250^ ATPase activity by the top candidate inhibitors (n=3 ± SD). **b**, Top candidate inhibitor binding curves, as determined by DSF (n=3 ± SD).

**Extended Data Fig. 2.**
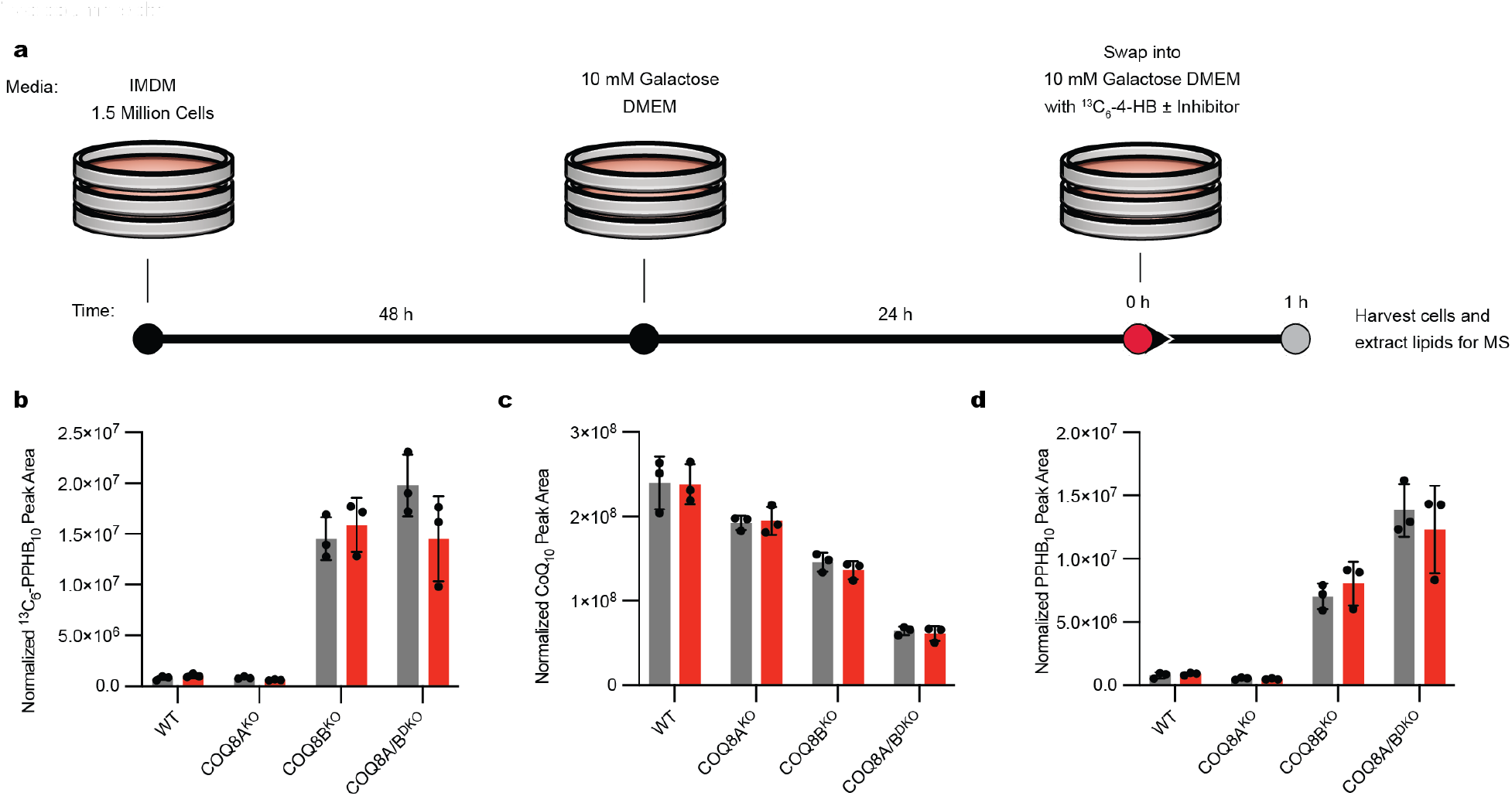
*De novo* CoQ_10_ biosynthesis measurements. **a**, Schematic outlining the experimental plan to measure *de novo* CoQ production. **b**, *De novo* production of ^13^C_6_-PPHB_10_ in WT, COQ8A^KO^, COQ8B^KO^, and COQ8A/B^DKO^ HAP1 cells after treatment with 10 μM ^13^C_6_-4-HB and either DMSO or 20 μM UNC-CA157 (grey=DMSO, red=UNC-CA157, n=3 ± SD). **c**, Unlabeled CoQ_10_ levels in WT, COQ8A^KO^, COQ8B^KO^, and COQ8A/B^DKO^ HAP1 cells after treatment with 10 μM ^13^C_6_-4-HB and either DMSO or 20 μM UNC-CA157 (grey=DMSO, red=UNC-CA157, n=3 ± SD). **d**, Unlabeled PPHB_10_ levels in WT, COQ8A^KO^, COQ8B^KO^, and COQ8A/B^DKO^ HAP1 cells after treatment with 10 μM ^13^C_6_-4-HB and either DMSO or 20 μM UNC-CA157 (grey=DMSO, red=UNC-CA157, n=3 ± SD).

**Extended Data Fig. 3.**
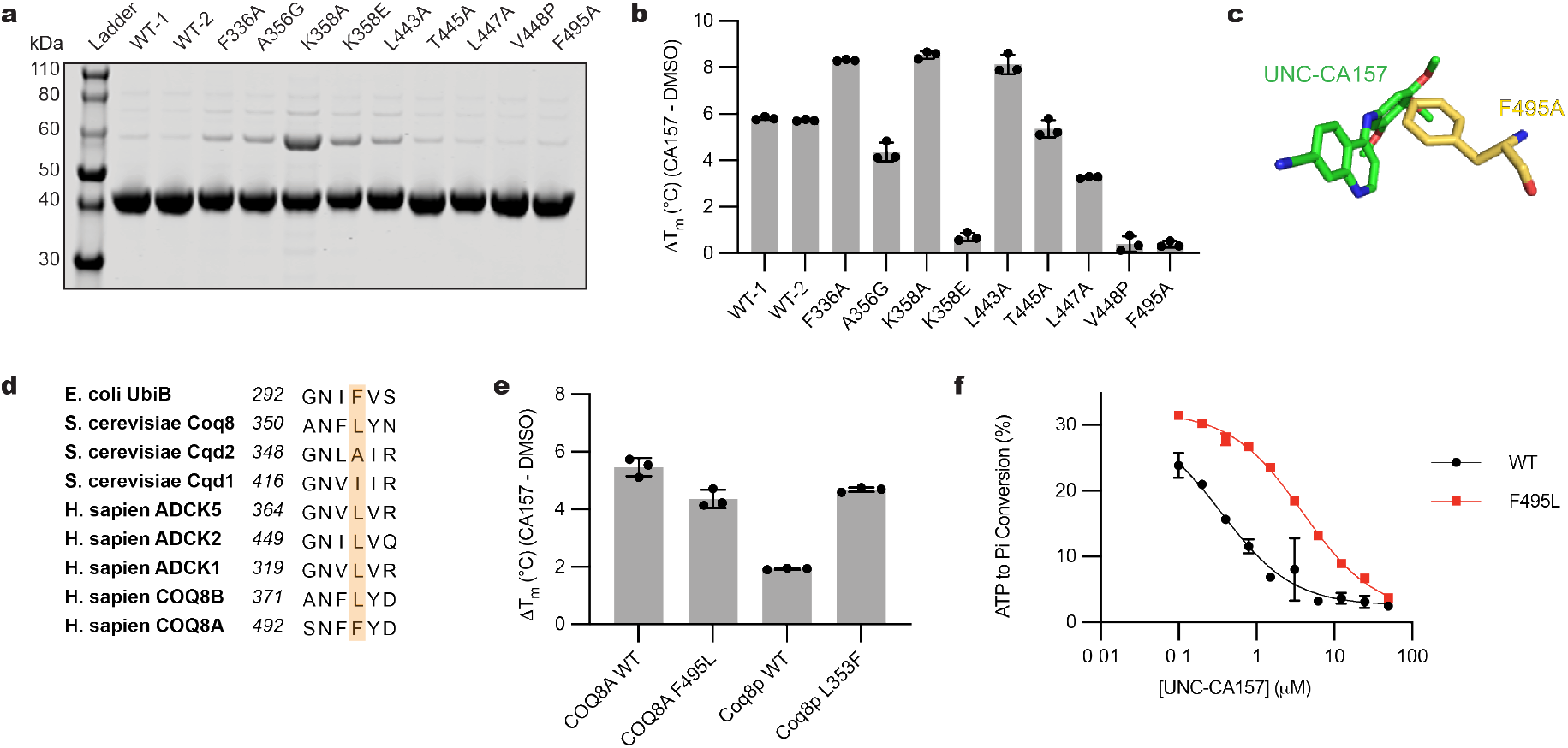
F495 is required for potent inhibition. **a**, SDS-PAGE analysis of COQ8A^NΔ250^ purifications. **b**, COQ8A^NΔ250^ WT and binding pocket mutant thermal stabilization by UNC-CA157, as determined by DSF (n=3 ± SD). **c**, Stick representation of UNC-CA157 and adjacent F495 residue. **d**, Conservation of F495 across the UbiB protein family. **e**, Thermal stabilization of COQ8A^NΔ250^ WT and F495L as well as Coq8p^NΔ41^ WT and L353F by UNC-CA157 (n=3 ± SD). **f**, Inhibition of COQ8A^NΔ250^ WT and F495L ATPase activity by UNC-CA157 (n=3 ± SD).

**Extended Data Fig. 4.**
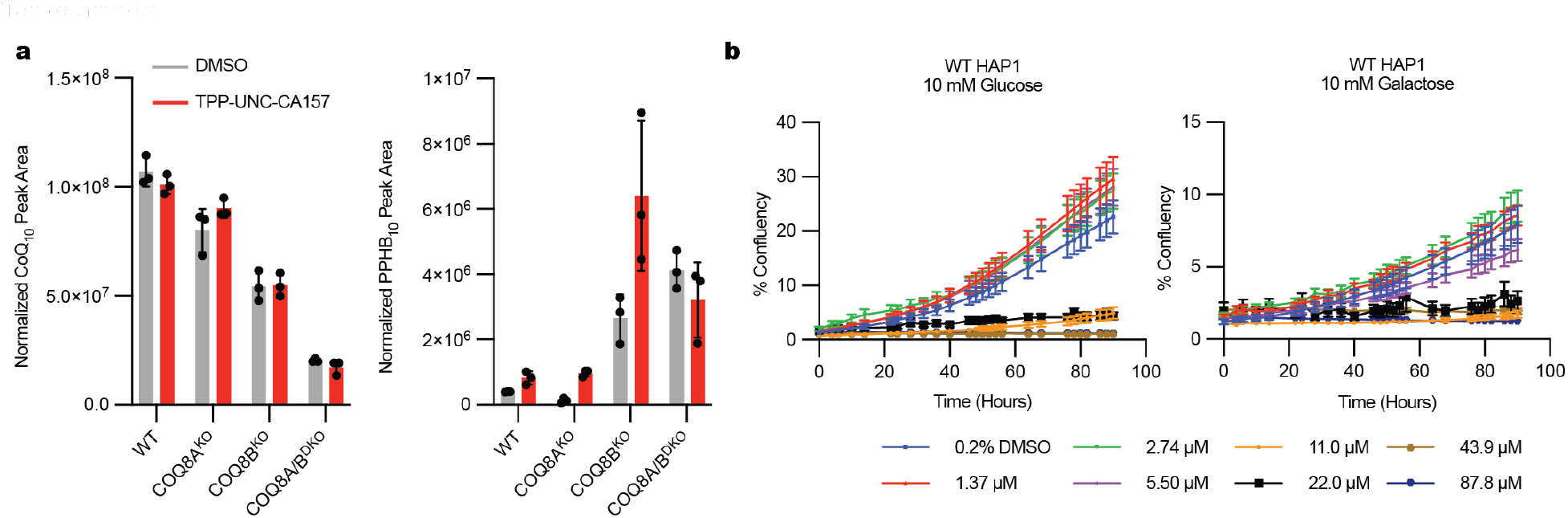
Results from *de novo* CoQ production with TPP-UNC-CA157 and inhibitor toxicity analysis. **a**, Unlabeled CoQ_10_ and PPHB_10_ levels in WT, COQ8A^KO^, COQ8B^KO^, and COQ8A/B^DKO^ HAP1 cells after treatment with 10 μM ^13^C_6_-4-HB and either DMSO or 17.6 μM TPP-UNC-CA157 (n=3 ± SD). **b**, Growth analysis of WT HAP1 cells in either 10 mM glucose or 10 mM galactose with indicated concentrations of TPP-UNC-CA157.

**Extended Data Fig. 5.**
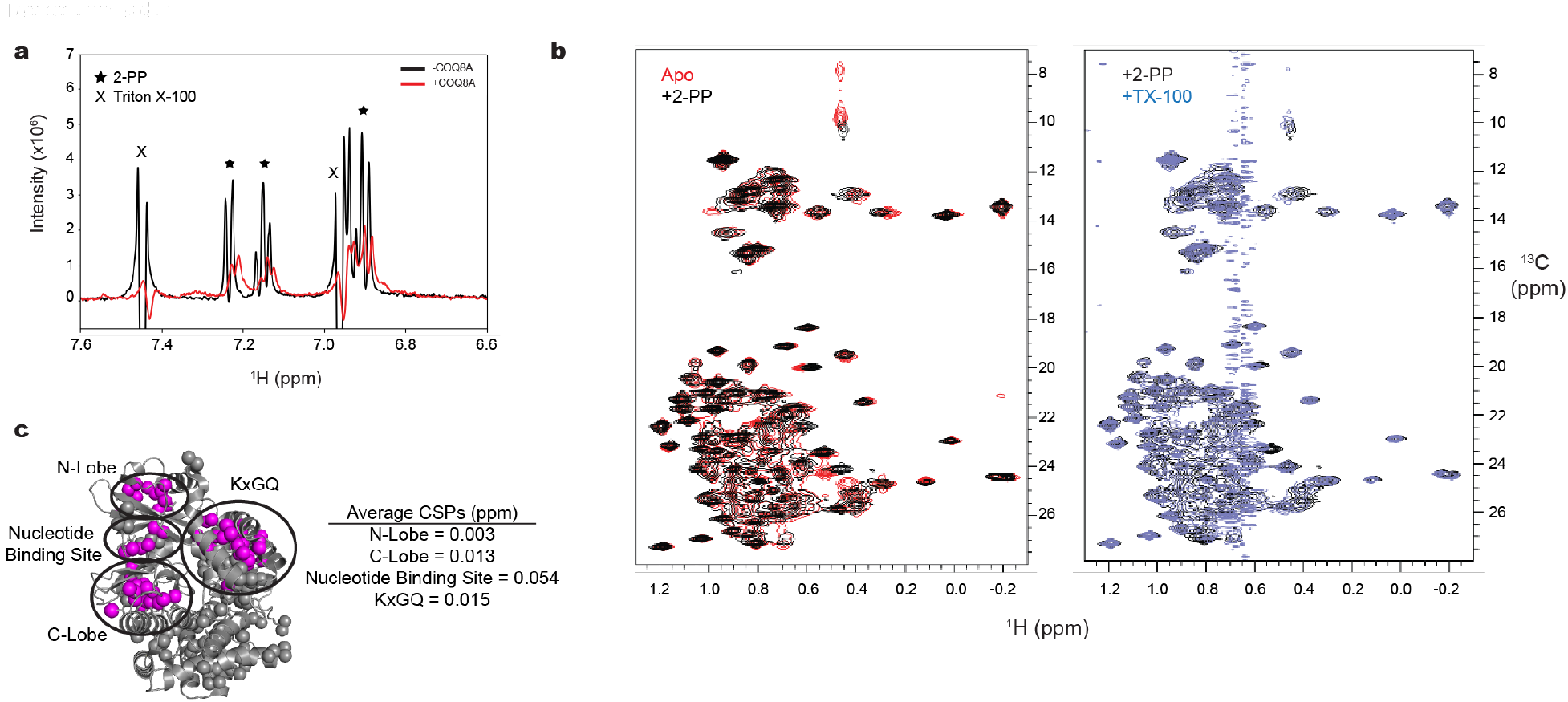
COQ8A NMR spectra and assigned ILV clusters. **a**, ^1^H NMR of 2-PP and TX-100 with and without COQ8A^NΔ250^. **b**, Overlaid ^1^H-^13^C HMQC spectra for COQ8A^NΔ250^ apo form vs. +1 mM 2-PP (left) and +1 mM 2-PP vs. +1 mM 2-PP, +1 mM TX-100 (right). **c**, Clustered ILV methyl assignments and average CSPs per cluster upon addition of 1 mM 2-PP.

**Extended Data Table 1.** Crystallography data collection and refinement statistics.

|  | **7UDP C2** | **7UDQ P1** |
| --- | --- | --- |
| **Wavelength** | 1.033 | 1.033 |
| **Resolution range** | 37.4 - 2.01 (2.082 - 2.01) | 38.72 - 1.9 (1.968 - 1.9) |
| **Space group** | C 1 2 1 | P 1 |
| **Unit cell** | 151.21 56.379 48.684 90<br>98.391 90 | 48.273 54.859 80.012<br>109.789 94.618 92.108 |
| **Total reflections** | 181981 (18308) | 211779 (21565) |
| **Unique reflections** | 27131 (2672) | 58805 (5808) |
| **Multiplicity** | 6.7 (6.8) | 3.6 (3.7) |
| **Completeness (%)** | 99.86 (99.93) | 96.99 (96.29) |
| **Mean I/sigma(I)** | 21.35 (1.01) | 9.38 (1.01) |
| **Wilson B-factor** | 65.59 | 41.87 |
| **R-merge** | 0.03374 (1.45) | 0.06329 (1.096) |
| **R-meas** | 0.03668 (1.569) | 0.07446 (1.281) |
| **R-pim** | 0.01422 (0.5943) | 0.03887 (0.6595) |
| **CC1/2** | 1 (0.511) | 0.997 (0.526) |
| **CC\*** | 1 (0.822) | 0.999 (0.83) |
| **Reflections used in refinement** | 27123 (2671) | 58710 (5807) |
| **Reflections used for R-free** | 2113 (210) | 2463 (254) |
| **R-work** | 0.2335 (0.3573) | 0.1993 (0.3104) |
| **R-free** | 0.2726 (0.3708) | 0.2383 (0.3254) |
| **CC(work)** | 0.960 (0.587) | 0.951 (0.756) |
| **CC(free)** | 0.930 (0.577) | 0.932 (0.692) |
| **Number of non-hydrogen atoms** | 3096 | 6354 |
| **macromolecules** | 3053 | 6036 |
| **ligands** | 42 | 86 |
| **solvent** | 18 | 266 |
| **Protein residues** | 378 | 747 |
| **RMS(bonds)** | 0.002 | 0.006 |
| **RMS(angles)** | 0.46 | 0.64 |
| **Ramachandran favored (%)** | 93.78 | 97.95 |
| **Ramachandran allowed (%)** | 6.22 | 1.91 |
| **Ramachandran outliers (%)** | 0.00 | 0.14 |
| **Rotamer outliers (%)** | 0.91 | 0.46 |
| **Clashscore** | 1.80 | 2.57 |
| **Average B-factor** | 87.97 | 55.62 |
| **macromolecules** | 88.28 | 55.96 |
| **ligands** | 66.95 | 41.00 |
| **solvent** | 64.56 | 50.89 |
| **Number of TLS groups** | 3 | 8 |

Statistics for the highest-resolution shell are shown in parentheses.

## References

1. Manning, G., Whyte, D. B., Martinez, R., Hunter, T. & Sudarsanam, S. The protein kinase complement of the human genome. Science 298, 1912–34 (2002).

2. Dar, A. C. & Shokat, K. M. The evolution of protein kinase inhibitors from antagonists to agonists of cellular signaling. Annu Rev Biochem 80, 769–95 (2011).

3. Cohen, P., Cross, D. & Jänne, P. A. Kinase drug discovery 20 years after imatinib: progress and future directions. Nat Rev Drug Discov 20, 551–569 (2021).

4. Attwood, M. M., Fabbro, D., Sokolov, A. V., Knapp, S. & Schiöth, H. B. Trends in kinase drug discovery: targets, indications and inhibitor design. Nat Rev Drug Discov 20, 839–861 (2021).

5. Kung, J. E. & Jura, N. Prospects for pharmacological targeting of pseudokinases. Nat Rev Drug Discov 18, 501–526 (2019).

6. Oprea, T. I. et al. Unexplored therapeutic opportunities in the human genome. Nat Rev Drug Discov 17, 317–332 (2018).

7. Ferguson, F. M. & Gray, N. S. Kinase inhibitors: the road ahead. Nat Rev Drug Discov 17, 353–377 (2018).

8. Leonard, C. J., Aravind, L. & Koonin, E. V. Novel families of putative protein kinases in bacteria and archaea: evolution of the “eukaryotic” protein kinase superfamily. Genome Res 8, 1038–47 (1998).

9. Pagliarini, D. J. et al. A mitochondrial protein compendium elucidates complex I disease biology. Cell 134, 112–23 (2008).

10. Lundquist, P. K., Davis, J. I. & Wijk, K. J. van. ABC1K atypical kinases in plants: filling the organellar kinase void. Trends Plant Sci 17, 546–55 (2012).

11. Lundquist, P. K. et al. Loss of plastoglobule kinases ABC1K1 and ABC1K3 causes conditional degreening, modified prenyl-lipids, and recruitment of the jasmonic acid pathway. Plant Cell 25, 1818–39 (2013).

12. Traschütz, A., et al. Clinico-Genetic, Imaging and Molecular Delineation of COQ8A-Ataxia: A Multicenter Study of 59 Patients. Ann Neurol 88, 251–263 (2020).

13. Mollet, J. et al. CABC1 gene mutations cause ubiquinone deficiency with cerebellar ataxia and seizures. Am J Hum Genetics 82, 623–30 (2008).

14. Lagier-Tourenne, C. et al. ADCK3, an ancestral kinase, is mutated in a form of recessive ataxia associated with coenzyme Q10 deficiency. Am J Hum Genetics 82, 661–72 (2008).

15. Stefely, J. A. et al. Cerebellar Ataxia and Coenzyme Q Deficiency through Loss of Unorthodox Kinase Activity. Mol Cell 63, 608–620 (2016).

16. Ashraf, S. et al. ADCK4 mutations promote steroid-resistant nephrotic syndrome through CoQ10 biosynthesis disruption. J Clin Invest 123, 5179–89 (2013).

17. Poon, W. W. et al. Identification of Escherichia coli ubiB, a gene required for the first monooxygenase step in ubiquinone biosynthesis. J Bacteriol 182, 5139–46 (2000).

18. Do, T. Q., Hsu, A. Y., Jonassen, T., Lee, P. T. & Clarke, C. F. A defect in coenzyme Q biosynthesis is responsible for the respiratory deficiency in Saccharomyces cerevisiae abc1 mutants. J Biol Chem 276, 18161–8 (2001).

19. Stefely, J. A. & Pagliarini, D. J. Biochemistry of Mitochondrial Coenzyme Q Biosynthesis. Trends Biochem Sci 42, 824–843 (2017).

20. Stefely, J. A. et al. Mitochondrial ADCK3 employs an atypical protein kinase-like fold to enable coenzyme Q biosynthesis. Mol Cell 57, 83–94 (2015).

21. Reidenbach, A. G. et al. Conserved Lipid and Small-Molecule Modulation of COQ8 Reveals Regulation of the Ancient Kinase-like UbiB Family. Cell Chem Biol 25, 154–165.e11 (2018).

22. Asquith, C. R. M., Murray, N. H. & Pagliarini, D. J. ADCK3/COQ8A: the choice target of the UbiB protein kinase-like family. Nat Rev Drug Discov 18, 815 (2019).

23. Asquith, C. R. M. et al. SGC-GAK-1: A Chemical Probe for Cyclin G Associated Kinase (GAK). J Med Chem 62, 2830–2836 (2019).

24. Wells, C. I. et al. The Kinase Chemogenomic Set (KCGS): An Open Science Resource for Kinase Vulnerability Identification. Int J Mol Sci 22, 566 (2021).

25. Robers, M. B. et al. Target engagement and drug residence time can be observed in living cells with BRET. Nat Commun 6, 10091 (2015).

26. Fonseca, L. V. et al. Mutations in COQ8B (ADCK4) found in patients with steroid-resistant nephrotic syndrome alter COQ8B function. Hum Mutat 39, 406–414 (2017).

27. Kanev, G. K. et al. The Landscape of Atypical and Eukaryotic Protein Kinases. Trends Pharmacol Sci 40, 818–832 (2019).

28. Zielonka, J. et al. Mitochondria-Targeted Triphenylphosphonium-Based Compounds: Syntheses, Mechanisms of Action, and Therapeutic and Diagnostic Applications. Chem Rev 117, 10043–10120 (2017).

29. Smith, R. A. J., Porteous, C. M., Gane, A. M. & Murphy, M. P. Delivery of bioactive molecules to mitochondria in vivo. Proc National Acad Sci 100, 5407–5412 (2003).

30. Trionnaire, S. L. et al. The synthesis and functional evaluation of a mitochondria-targeted hydrogen sulfide donor, (10-oxo-10-(4-(3-thioxo-3 H −1,2-dithiol-5-yl)phenoxy)decyl)triphenylphosphonium bromide (AP39). Medchemcomm 5, 728–736 (2014).

31. Trnka, J., Elkalaf, M. & Anděl, M. Lipophilic Triphenylphosphonium Cations Inhibit Mitochondrial Electron Transport Chain and Induce Mitochondrial Proton Leak. Plos One 10, e0121837 (2015).

32. Reily, C. et al. Mitochondrially targeted compounds and their impact on cellular bioenergetics. Redox Biol 1, 86–93 (2013).

33. Huang, R. & Leung, I. K. H. Protein–Small Molecule Interactions by WaterLOGSY. Methods Enzymol 615, 477–500 (2018).

34. Engen, J. R., Botzanowski, T., Peterle, D., Georgescauld, F. & Wales, T. E. Developments in Hydrogen/Deuterium Exchange Mass Spectrometry. Anal Chem 93, 567–582 (2021).

35. Serafim, R. A. M., Elkins, J. M., Zuercher, W. J., Laufer, S. A. & Gehringer, M. Chemical Probes for Understudied Kinases: Challenges and Opportunities. J Med Chem 65, 1132–1170 (2022).

36. Kung, J. E. & Jura, N. Structural Basis for the Non-catalytic Functions of Protein Kinases. Structure 24, 7–24 (2016).

37. Miles, M. V. The uptake and distribution of coenzyme Q(10). Mitochondrion 7, S72–S77 (2007).

38. He, C. H., Xie, L. X., Allan, C. M., Tran, U. C. & Clarke, C. F. Coenzyme Q supplementation or over-expression of the yeast Coq8 putative kinase stabilizes multi-subunit Coq polypeptide complexes in yeast coq null mutants. Biochimica Et Biophysica Acta Bba - Mol Cell Biology Lipids 1841, 630–44 (2014).

39. Xie, L. X. et al. Overexpression of the Coq8 kinase in Saccharomyces cerevisiae coq null mutants allows for accumulation of diagnostic intermediates of the coenzyme Q6 biosynthetic pathway. J Biol Chem 287, 23571–81 (2012).

40. Larsen, P. L. & Clarke, C. F. Extension of life-span in Caenorhabditis elegans by a diet lacking coenzyme Q. Science 295, 120–3 (2002).

41. Liu, X. et al. Evolutionary conservation of the clk-1-dependent mechanism of longevity: loss of mclk1 increases cellular fitness and lifespan in mice. Gene Dev 19, 2424–2434 (2005).

42. Doll, S. et al. FSP1 is a glutathione-independent ferroptosis suppressor. Nature 575, 693–698 (2019).

43. Bersuker, K. et al. The CoQ oxidoreductase FSP1 acts parallel to GPX4 to inhibit ferroptosis. Nature 575, 688–692 (2019).

44. Forsman, U., Sjöberg, M., Turunen, M. & Sindelar, P. J. 4-Nitrobenzoate inhibits coenzyme Q biosynthesis in mammalian cell cultures. Nat Chem Biol 6, 515–517 (2010).

45. Wang, Y. et al. The Anti-neurodegeneration Drug Clioquinol Inhibits the Aging-associated Protein CLK-1*. J Biol Chem 284, 314–323 (2009).

46. Fernández-del-Río, L. et al. Regulation of hepatic coenzyme Q biosynthesis by dietary omega-3 polyunsaturated fatty acids. Redox Biol 102061 (2021) doi:10.1016/j.redox.2021.102061.

47. Kelso, G. F. et al. Selective Targeting of a Redox-active Ubiquinone to Mitochondria within Cells. J Biol Chem 276, 4588–4596 (2001).

48. Kulkarni, C. A. et al. A Novel Triphenylphosphonium Carrier to Target Mitochondria without Uncoupling Oxidative Phosphorylation. J Med Chem 64, 662–676 (2021).

49. Kemmerer, Z. A. et al. UbiB proteins regulate cellular CoQ distribution in Saccharomyces cerevisiae. Nat Commun 12, 4769 (2021).

## Methods references

1. Stefely, J. A. et al. Mitochondrial ADCK3 employs an atypical protein kinase-like fold to enable coenzyme Q biosynthesis. Mol Cell 57, 83–94 (2015).

2. Fox, B. G. & Blommel, P. G. Autoinduction of Protein Expression. Curr Protoc Protein Sci 56, 5.23.1–5.23.18 (2009).

3. Asquith, C. R. M. et al. Identification and Optimization of 4-Anilinoquinolines as Inhibitors of Cyclin G Associated Kinase. Chemmedchem 13, 48–66 (2017).

4. Asquith, C. R. M. et al. Design of a Cyclin G Associated Kinase (GAK)/Epidermal Growth Factor Receptor (EGFR) Inhibitor Set to Interrogate the Relationship of EGFR and GAK in Chordoma. J Med Chem 62, 4772–4778 (2019).

5. Asquith, C. R. M. et al. Design and Analysis of the 4-Anilinoquin(az)oline Kinase Inhibition Profiles of GAK/SLK/STK10 Using Quantitative Structure-Activity Relationships. Chemmedchem 15, 26–49 (2019).

6. Asquith, C. R. M. et al. SGC-GAK-1: A Chemical Probe for Cyclin G Associated Kinase (GAK). J Med Chem 62, 2830–2836 (2019).

7. Asquith, C. R. M. et al. Targeting an EGFR Water Network with 4-Anilinoquin(az)oline Inhibitors for Chordoma. Chemmedchem 14, 1693–1700 (2019).

8. Asquith, C. R. M., Treiber, D. K. & Zuercher, W. J. Utilizing comprehensive and mini-kinome panels to optimize the selectivity of quinoline inhibitors for cyclin G associated kinase (GAK). Bioorg Med Chem Lett 29, 1727–1731 (2019).

9. Niesen, F. H., Berglund, H. & Vedadi, M. The use of differential scanning fluorimetry to detect ligand interactions that promote protein stability. Nat Protoc 2, 2212–21 (2007).

10. Reidenbach, A. G. et al. Conserved Lipid and Small-Molecule Modulation of COQ8 Reveals Regulation of the Ancient Kinase-like UbiB Family. Cell Chem Biol 25, 154–165.e11 (2018).

11. Kueng, W., Silber, E. & Eppenberger, U. Quantification of cells cultured on 96-well plates. Anal Biochem 182, 16–19 (1989).

12. Stepanov, S. et al. JBluIce-EPICS control system for macromolecular crystallography. Acta Crystallogr Sect D Biological Crystallogr 67, 176–88 (2011).

13. Vonrhein, C. et al. Data processing and analysis with the autoPROC toolbox. Acta Crystallogr Sect D Biological Crystallogr 67, 293–302 (2011).

14. Kabsch, W. XDS. Acta Crystallogr Sect D Biological Crystallogr 66, 125–32 (2010).

15. McCoy, A. J., et al. Phaser crystallographic software. J Appl Crystallogr 40, 658–674 (2007).

16. Moriarty, N. W., Grosse-Kunstleve, R. W. & Adams, P. D. electronic Ligand Builder and Optimization Workbench (eLBOW): a tool for ligand coordinate and restraint generation. Acta Crystallogr Sect D Biological Crystallogr 65, 1074–80 (2009).

17. Emsley, P., Lohkamp, B., Scott, W. G. & Cowtan, K. Features and development of Coot. Acta Crystallogr Sect D Biological Crystallogr 66, 486–501 (2010).

18. Afonine, P. V. et al. Towards automated crystallographic structure refinement with phenix.refine. Acta Crystallogr Sect D Biological Crystallogr 68, 352–367 (2012).

19. Williams, C. J. et al. MolProbity: More and better reference data for improved all-atom structure validation. Protein Sci 27, 293–315 (2018).

20. Yang, H. et al. Automated and accurate deposition of structures solved by X-ray diffraction to the Protein Data Bank. Acta Crystallogr Sect D Biological Crystallogr 60, 1833–1839 (2004).

21. Morin, A. et al. Collaboration gets the most out of software. Elife 2, e01456 (2013).

22. Stefely, J. A. et al. Cerebellar Ataxia and Coenzyme Q Deficiency through Loss of Unorthodox Kinase Activity. Mol Cell 63, 608–620 (2016).

23. Trionnaire, S. L. et al. The synthesis and functional evaluation of a mitochondria-targeted hydrogen sulfide donor, (10-oxo-10-(4-(3-thioxo-3H-1,2-dithiol-5-yl)phenoxy)decyl)triphenylphosphonium bromide (AP39). Med Chem Commun 5, 728–736 (2014).

24. Delaglio, F. et al. NMRPipe: A multidimensional spectral processing system based on UNIX pipes. J Biomol Nmr 6, 277–93 (1995).

25. Helmus, J. J. & Jaroniec, C. P. Nmrglue: an open source Python package for the analysis of multidimensional NMR data. J Biomol Nmr 55, 355–367 (2013).

26. Cai, M. et al. An efficient and cost-effective isotope labeling protocol for proteins expressed in Escherichia coli. J Biomol Nmr 11, 97–102 (1998).

27. Lichtenecker, R. J. et al. Independent valine and leucine isotope labeling in Escherichia coli protein overexpression systems. J Biomol Nmr 57, 205–9 (2013).

28. Tugarinov, V., Kanelis, V. & Kay, L. E. Isotope labeling strategies for the study of high-molecular-weight proteins by solution NMR spectroscopy. Nat Protoc 1, 749–754 (2006).

29. Tugarinov, V., Kay, L. E., Ibraghimov, I. & Orekhov, V. Y. High-resolution four-dimensional 1H-13C NOE spectroscopy using methyl-TROSY, sparse data acquisition, and multidimensional decomposition. J Am Chem Soc 127, 2767–75 (2005).

30. Ying, J., Delaglio, F., Torchia, D. A. & Bax, A. Sparse multidimensional iterative lineshape-enhanced (SMILE) reconstruction of both non-uniformly sampled and conventional NMR data. J Biomol Nmr 68, 101–118 (2017).

31. Vranken, W. F. et al. The CCPN data model for NMR spectroscopy: development of a software pipeline. Proteins 59, 687–96 (2005).

32. Gorman, S. D., Sahu, D., O’Rourke, K. F. & Boehr, D. D. Assigning methyl resonances for protein solution-state NMR studies. Methods 148, 88–99 (2018).

33. Pabon, J. P. R., Kochert, B. A., Liu, Y.-H., Richardson, D. D. & Weis, D. D. Protein A does not induce allosteric structural changes in an IgG1 antibody during binding. J Pharm Sci 110, 2355–2361 (2021).

34. Glasoe, P. K. & Long, F. A. USE OF GLASS ELECTRODES TO MEASURE ACIDITIES IN DEUTERIUM OXIDE 1,2. J Phys Chem 64, 188–190 (1960).

35. Hageman, T. S. & Weis, D. D. Reliable Identification of Significant Differences in Differential Hydrogen Exchange-Mass Spectrometry Measurements Using a Hybrid Significance Testing Approach. Anal Chem 91, 8008–8016 (2019).

