## Supplementary material for "Small Molecule Modulation of the Archetypal UbiB protein COQ8": HDX-MS Supplement

### Supplementary HDX-MS Information

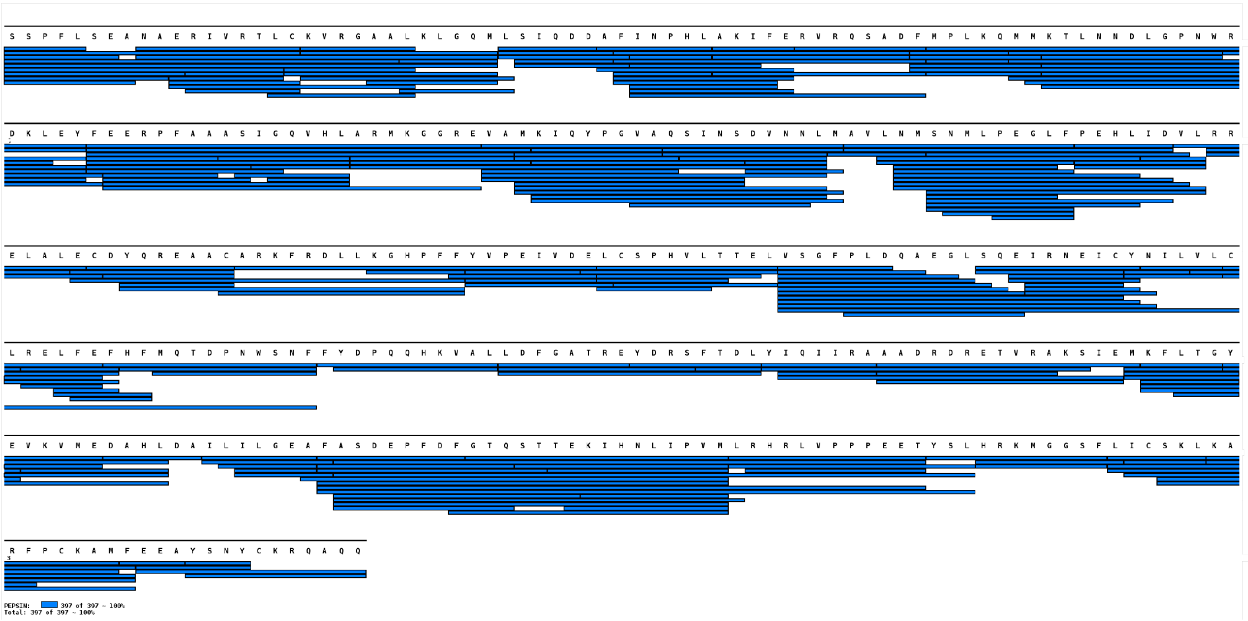


**Figure 1.** COQ8A$^{N\Delta 250}$ peptic peptides coverage map. Coverage map was made using MSTools[1].

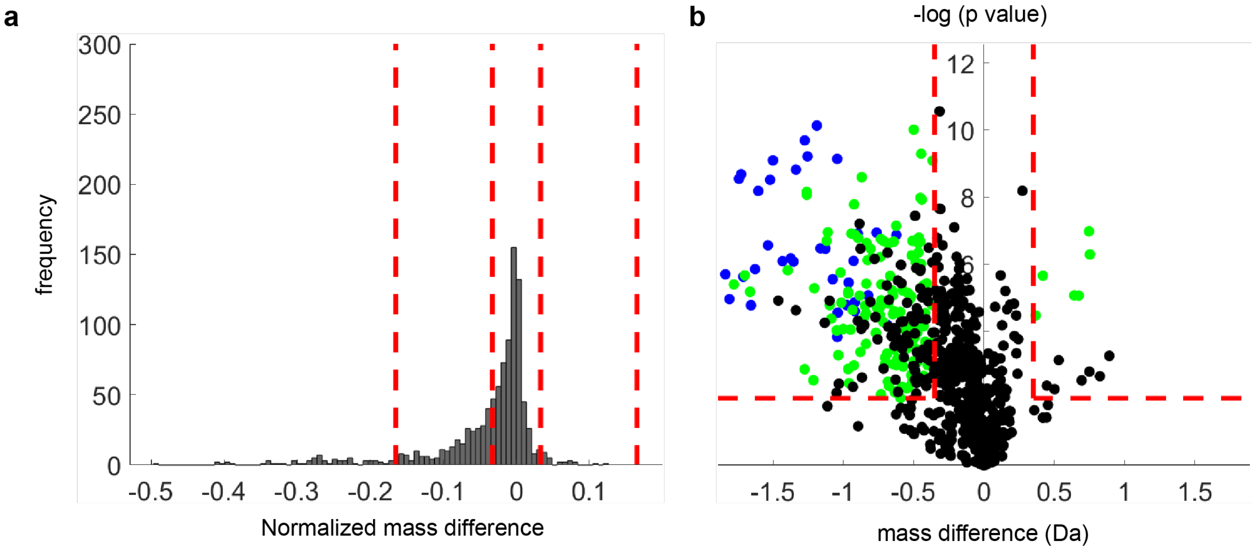


**Figure 2.** HDX-MS statistical analyses for identification of significant differences. (**a**) Binning of normalized HDX differences in deuterium uptake for categorization into strong, moderate, and negligible differences. (**b**) Volcano plot for evaluation of statistically different peptides and the magnitude of the effects (see methods section). Negligible (black), moderate (green), and strong (blue) effects as revealed by k-mean clustering of the data.

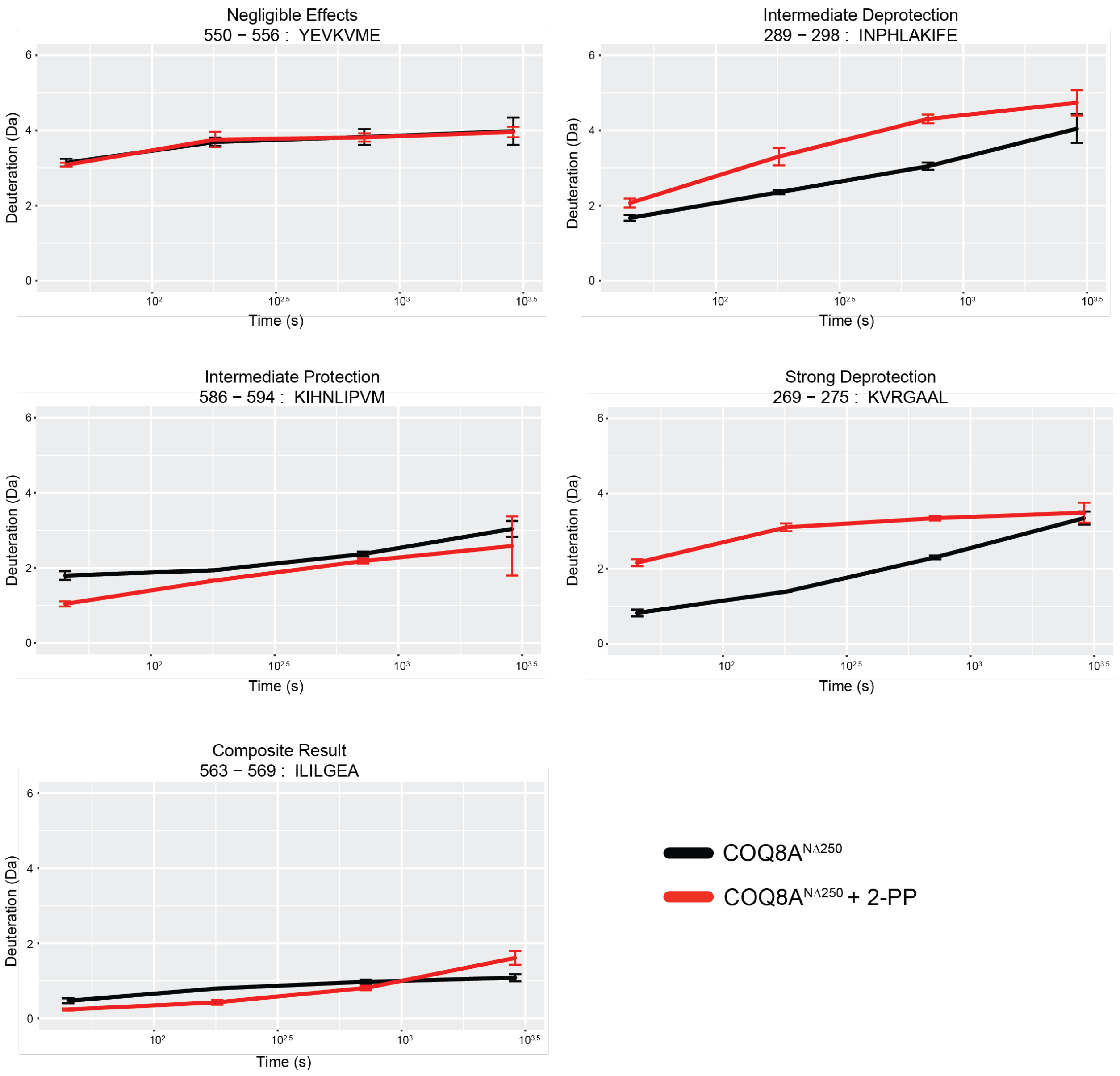


**Figure 3.** Representative uptake plots for different peptide classifications showing the effect of 2-PP binding to COQ8A$^{N\Delta250}$. Error bars represent the standard deviation of the measurements with n=3.

#### References

1. Kavan D, Man P 2011. MSTools—Web based application for visualization and presentation of HXMS data. International Journal of Mass Spectrometry 302(1):53-58.
