## Supplementary material for "Small Molecule Modulation of the Archetypal UbiB protein COQ8": Inhibitor Supplement

### Supplementary Inhibitor Information

#### Compounds from Figure 1a

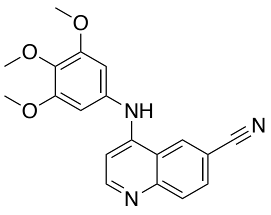

UNC-CA171

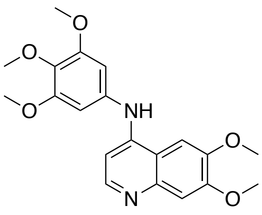

UNC-CA75

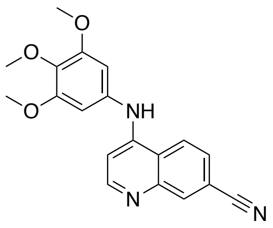

UNC-CA157

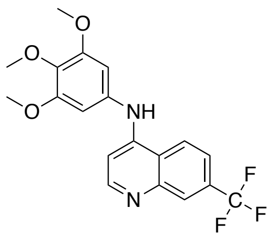

UNC-CA80

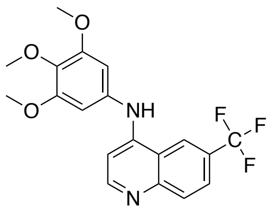

UNC-CA62

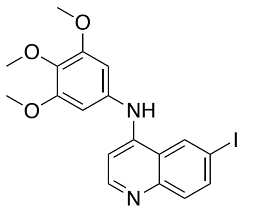

UNC-CA92

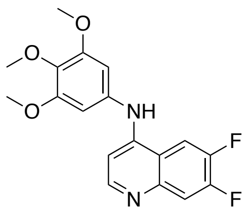

UNC-CA327

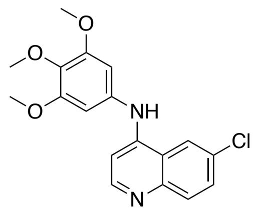

UNC-CA169

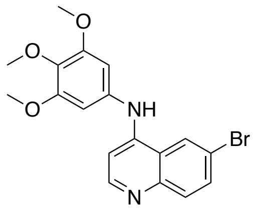

UNC-CA93

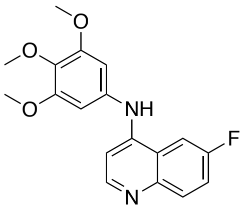

UNC-CA82

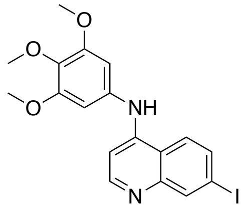

UNC-CA94

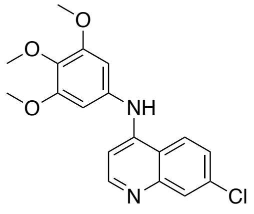

UNC-CA173

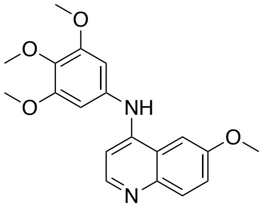

UNC-CA81

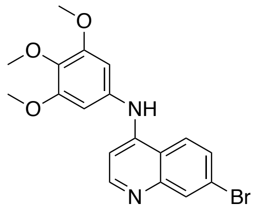

UNC-CA93.5

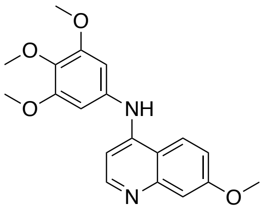

UNC-CA162

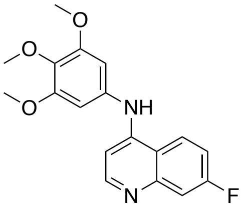

UNC-CA163

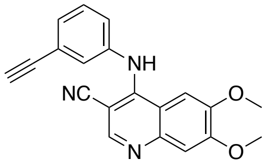

UNC-CA252

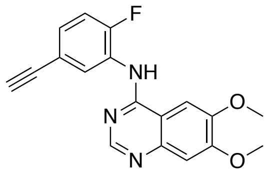

UNC-CA360

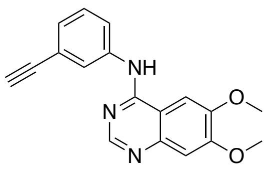

UNC-CA176

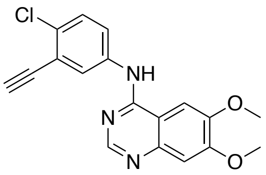

UNC-CA359

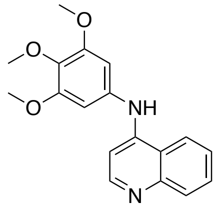

UNC-CA73

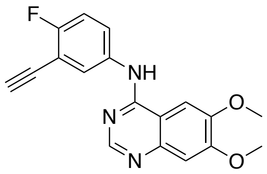

UNC-CA358

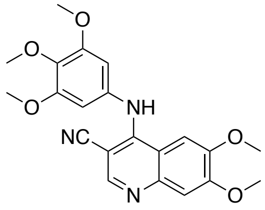

UNC-CA249

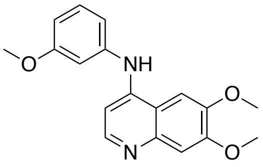

UNC-CA128

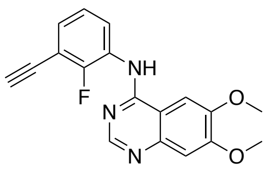

UNC-CA384

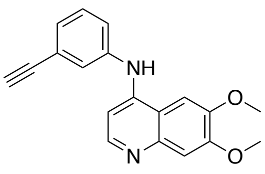

UNC-CA156

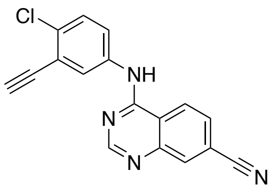

UNC-CA331

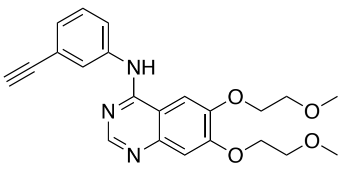

Erlotinib

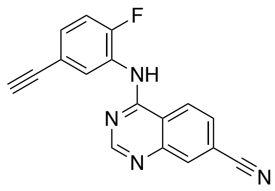

UNC-CA332

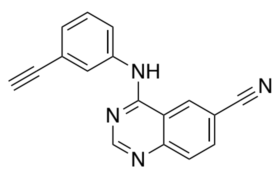

UNC-CA207

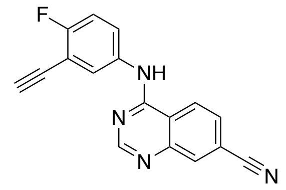

UNC-CA330

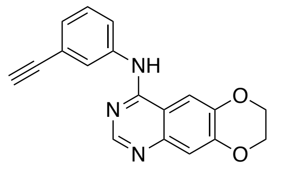

UNC-CA309

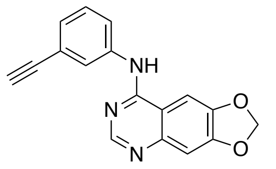

UNC-CA310

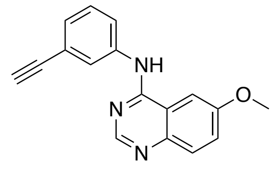

UNC-CA209

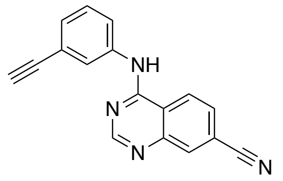

UNC-CA271

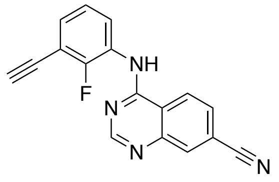

UNC-CA386

**Inhibition data from Extended Data Figure 1a**

| Compound | $IC_{50}$ (μM) | 95% Confidence Interval (μM) |
|---|---|---|
| UNC-CA171 | 0.69 | ± 0.11 |
| UNC-CA75 | 0.71 | ± 0.10 |
| UNC-CA157 | 0.47 | ± 0.076 |
| UNC-CA80 | 0.49 | ± 0.10 |
| UNC-CA92 | 1.5 | ± 0.21 |
| UNC-CA327 | 1.1 | ± 0.23 |
| UNC-CA169 | 1.2 | ± 0.17 |
| UNC-CA62 | 1.7 | ± 0.38 |

**DSF binding data from Extended Data Figure 1b**

| Compound | $K_{d,app}$ (μM) | 95% Confidence Interval (μM) |
|---|---|---|
| UNC-CA171 | 26 | ± 9.1 |
| UNC-CA75 | 15 | ± 2.7 |
| UNC-CA157 | 23 | ± 9.2 |
| UNC-CA80 | 7.6 | ± 4.3 |
| UNC-CA92 | 21 | ± 6.5 |
| UNC-CA327 | 27 | ± 6.6 |
| UNC-CA169 | 20 | ± 7.5 |
| UNC-CA62 | 26 | ± 9.9 |

**NanoBRET apparent live cell affinity data**

| **Compound** | **COQ8A $IC_{50}$ (nM)** | **COQ8B $IC_{50}$ (nM)** |
|---|---|---|
| **PD173955** | **1080 ± 70** | **270 ± 30** |
| **UNC-CA157** | **580 ± 40** | **> 10,000** |
| **UNC-CA171** | **730 ± 40** | **> 10,000** |
| **UNC-CA75** | **1410 ± 90** | **> 10,000** |
| **UNC-CA81** | **7000 ± 3000** | **> 10,000** |
| **UNC-CA162** | **3600 ± 500** | **> 10,000** |

**NanoBRET assay validation data**

**a**

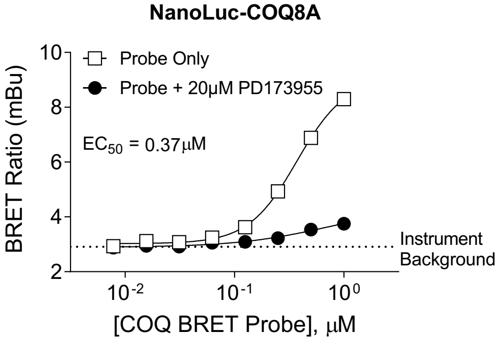


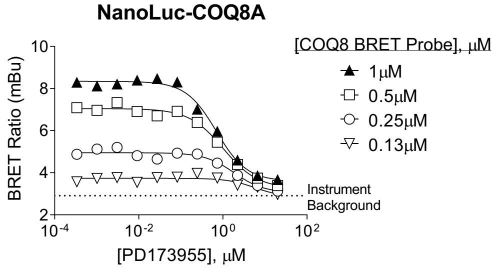


| | 1µM | 0.5µM | 0.25µM | 0.13µM |
|---|---|---|---|---|
| IC50 | 0.70 | 1.0 | 1.9 | 5.7 |

**b**

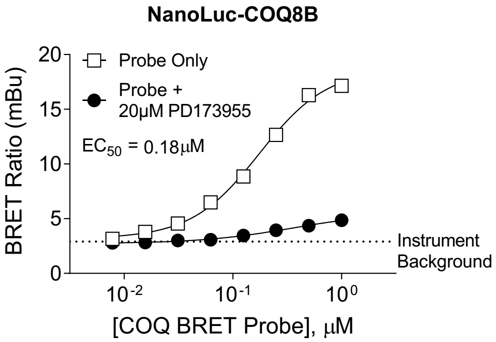


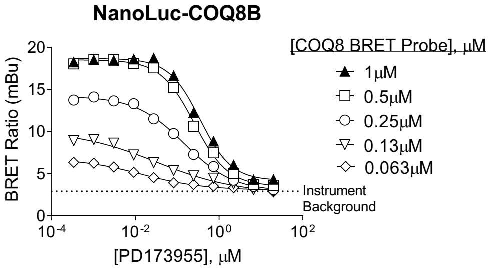


| | 1µM | 0.5µM | 0.25µM | 0.13µM | 0.063µM |
|---|---|---|---|---|---|
| IC50 | 0.35 | 0.25 | 0.15 | 0.032 | 0.014 |

**Characterization of BRET assay for COQ8A and COQ8B in live cells.** Characterization of BRET between (A) NanoLuc-COQ8A and (B) NanoLuc-COQ8B and the COQ8 BRET probe. Probe titrations (left) produced specific BRET as demonstrated via competition with 20µM PD173955. Individual data points are the mean of technical duplicates (n=1). PD173955 competed the BRET signal in a dose-responsive manner (right). Individual data points are technical singlicates (n=1). The probe concentrations used for subsequent affinity measurements for test compounds are highlighted in solid black triangles (right).
