## Supplementary material for "Small Molecule Modulation of the Archetypal UbiB protein COQ8": NMR Supplement

### Supplementary NMR Information

#### Protein-Observed NMR

NMR studies employed the COQ8A$^{N\Delta 250}$ construct lacking the N-terminal 250 residues. The construct is 46 kDa, near the upper limit of the molecular weight range where a backbone-directed approach is viable. As such, we opted for a methyl-labeling approach more amenable to large proteins[1,2]. In this scheme, the terminal methyl groups of Ile (δ1), Leu (δ1/δ2), and Val (γ1/γ2) are selectively $^{13}C$, $^{1}H$ labeled in a perdeuterated background. COQ8A$^{N\Delta 250}$ contains 21 Ile, 20 Val, and 43 Leu residues, providing a significant number of probes to detect binding. The chosen labeling scheme yields one peak per Ile and 2 peaks per Val and Leu, leading to an expected total of 147 peaks in the spectrum.

A $^{1}H$-$^{13}C$ HMQC spectrum was collected for apo COQ8A$^{N\Delta 250}$ (Fig. 1a). The spectrum showed well-dispersed chemical shifts, indicating that the protein is stably folded. Ile residues fall in a distinct region of the spectrum (~11-15 ppm in the $^{13}C$ dimension); 21/21 expected Ile peaks were observed. Leu and Val chemical shift ranges partially overlap, but 104/126 expected peaks were observed in the L/V region. In total, 85% of expected peaks were observed. Observation of fewer than the expected number of peaks is likely due to two factors. First, the Leu region of the spectrum (around ~25 ppm in the $^{13}C$ dimension) is very crowded and may contain more peaks than can be uniquely identified by visual inspection. Second, there may be intermediate timescale (~μs-ms) internal dynamics present in the protein causing peak broadening. Several peaks are visibly broadened in the apo spectrum, supporting the notion that some peaks are likely missing due to peak broadening.

Addition of 1 mM 2-PP to COQ8A$^{N\Delta 250}$ induced chemical shift perturbations (CSPs), again confirming the binding interaction (Fig. 1). Due to crowding, not every peak can be uniquely identified in both apo and +2-PP spectra, but 33% of peaks picked in both spectra undergo CSPs at least 1.5x the 10% trimmed mean CSP. This could indicate that 2-PP is exerting an allosteric effect on the protein structure. Several resonances are reduced in intensity upon addition of 2-PP as well. Broadening could be caused by binding and unbinding of 2-PP or could indicate that 2-PP otherwise affects or induces internal dynamics in COQ8A$^{N\Delta250}$. Further addition of 1 mM TX-100 caused CSPs, although fewer than 2-PP (Fig. 1). TX-100 also induced precipitation of COQ8A$^{N\Delta250}$, likely due to the high concentrations of TX-100 and protein used in the experiment. Further NMR studies with TX-100 were not pursued due to the observed precipitation. Overall, these results showed that we can produce methyl-labeled samples of folded COQ8A$^{N\Delta250}$ and that 2-PP binding to COQ8A$^{N\Delta250}$ can be observed by solution NMR.

#### Resonance assignment of ILV methyl groups in COQ8A$^{N\Delta250}$

Since 2-PP induced CSPs in COQ8A$^{N\Delta250}$, the binding site can be mapped once peaks assignments are obtained. Ideally, peaks that shift upon ligand addition will cluster to a certain location on the structure, revealing the binding site. NOESY-based approaches were used to assign methyl resonances in COQ8A$^{N\Delta250}$. This approach entails recording a NOESY spectrum of an ILV-labeled protein, which correlates a given methyl resonance with the resonances for other methyl groups that are physically close in space[2,3]. Using this data, graph matching algorithms can match the observed methyl connectivity network with the expected network from the crystal structure[4,5]. It is not typically possible to assign all methyl resonances from the NOESY data alone, but in benchmarking the graph matching algorithms achieved assignment completeness varying from 37%-89% depending on the target[4,5].

A 4D HCCH HMQC-NOESY-HMQC spectrum[3] was recorded for apo COQ8A$^{N\Delta250}$. The 4D approach was employed due to the crowded nature of the L/V region in the COQ8A$^{N\Delta250}$ spectrum. An example 2D plane from the 4D experiment is shown in Figure 2. Manual peak picking of the 4D spectrum yielded a total of 223 NOEs for 125 peaks in the 2D spectrum. MAGMA software[4] was used as the first attempt to assign COQ8A$^{N\Delta250}$ from the 4D NOESY data alone. The 4ped structure of apo COQ8A was used to generate the structure graph. Unfortunately, the algorithm could not converge using only the 4D NOESY data, likely due to the fairly low number of total NOEs observed. This may occur when internal protein dynamics impede magnetization transfer via the NOE. Indeed, many peaks are visibly broadened and less than the expected number of peaks are present in the spectrum. Both observations support the presence of internal dynamics in COQ8A$^{N\Delta250}$.

Several avenues were taken to reduce ambiguity in the NOESY data and increase assignment completeness. First, a sample was produced where only Val, but not Leu residues were labeled (Fig. 3). This allowed for discrimination between L/V residue types. Second, a sample was produced where both γ (for Val) or δ (for Leu) methyl groups were $^{13}C$ labeled, as opposed to where only one of two is labeled in the previous scheme. A 3D NOESY-HMQC spectrum with a shorter mixing time was collected for this sample, allowing identification of geminal methyl groups (Fig. 4). In most cases, only one NOE is observed per peak and that NOE corresponds to the geminally-connected methyl group. The NOE can be confirmed to have arisen from the geminal methyl group by seeing that it is absent from the 4D NOESY, where only one of two methyls are labeled per L/V. Providing L/V residue type discrimination and L/V geminal pairing greatly reduces the complexity of the data graph for the graph matching algorithms, but assignments could still not be attained.

Subsequently, an additional 3D NOESY dataset was collected for AMPPNP-bound COQ8A$^{N\Delta250}$. We reasoned that binding of a ligand may help stabilize the protein and reduce some of the internal dynamics, but spectra of AMPPNP-bound COQ8A$^{N\Delta250}$ revealed a similar amount of line-broadening. Also, eliminating one chemical shift evolution period from the pulse sequence by going from 4D to 3D data collected should also increase signal-to-noise in the resulting spectrum, of course at the expense of resolution. The 3D NOESY spectrum of COQ8A$^{N\Delta250}$ in the presence of 2 mM AMPPNP did reveal a greater number of NOEs than the 4D spectrum. In total, 317 non-diagonal NOEs were picked. Unfortunately, having one fewer chemical shift dimension meant that some NOEs could not be unambiguously assigned.

NOESY data was also supplemented with mutational data to fix assignments. Mutants used for assignment include V448A, L447A, L506A, V344A, I283L, I471L, I359L, I296L I525L, I563L, I591L,

I368L, V378A, I587L, and I395L (Main text Fig. 5c). Despite the additional data, all attempts at using MAGMA[4] or MAUS[5] to obtain further assignments were unsuccessful.

Despite difficulty in obtaining a high degree assignment completeness from algorithm-based approaches, manual analysis of NOESY patterns in conjunction with mutational data has been helpful in obtaining some assignments (Main text Fig. 5d). Mutation sites were chosen in important regions of the protein (eg nucleotide binding site) or because the given residue would be predicted to have a large number of NOEs based on the structure. In particular, the I471L mutation allowed for the assignment of a cluster of residues at the interface of β7/β8 with the αE and αF helices in the C-lobe. Analysis of NOESY patterns from I359 allowed for nearly complete assignments in the N-lobe while assignments in the KxGQ domain were enabled by hard assignments via mutation for I296, V378, I395, and I368. Using the mutational data, a total of 57 peaks in the 2D spectrum could be confidently assigned, corresponding to 39 residues with at least one methyl group assigned.

Spectra of the I525L, I563L, I587L, and I591L mutants did not show an obvious peak disappearance in the Ile region. It is likely then that these residues undergo ms timescale dynamics leading to broadening and peak disappearance. I525, I563, I587 and I591 are all found in the C-lobe. As noted above, fewer than the expected number of peaks are observed in the 2D spectrum. Thus, it is possible that missing peaks originate from residues in the C-lobe. The absence of peaks for these residues could also explain why the structure-based approaches have failed in yielding assignments; if large portions of the structure are not represented in the NMR data, the data graph and structure graph will not match well.

In summary, a combination of NOESY and mutational data allowed for the assignment of at least one methyl group in 39 residues in COQ8A$^{N\Delta250}$. Assigned residues cluster in the N-lobe (cluster 1), nucleotide binding site (cluster 2) KxGQ domain (cluster 3), and part of the C-lobe (cluster 4) (Main text Extended Data Fig. 5c). Mutated residues in the Cα 1-4 section of the C-lobe seem to be missing from the spectrum, indicating that there may be ms timescale dynamics in the C-lobe leading to peak disappearance.

Absence of data for residues in the C-lobe may be hindering further assignments by algorithm-based approaches.

#### 2-PP induced CSPs in COQ8A$^{N\Delta250}$

Surprisingly, residues in all assigned clusters other than the N-lobe undergo significant CSPs upon 2-PP binding. CSPs for assigned residues are plotted on the apo structure of COQ8A in Main text Fig. 5f. Methyl groups are not currently stereospecifically assigned, so an average CSP was calculated from both geminal methyl groups and that average value was plotted. For each cluster (clusters 1-4), average CSP values were calculated for assigned residues plus peaks for unassigned residues that have NOEs to assigned residues in a given cluster. The N-lobe (cluster 1) was essentially unaffected by 2-PP binding, with an average CSP of 0.003 ppm. Large CSPs were observed for residues V448, L506, and V344 in the nucleotide binding sites (cluster 2), where the average CSP is 0.054 ppm. In fact, L506 and V344 experience some of the largest CSPs of all peaks in the spectrum. In cluster 4, adjacent to the nucleotide binding site, the average CSP was 0.013. Similarly, the KxGQ domain (cluster 3) had an average CSP of 0.015.

Identifying the 2-PP binding site from the CSP data is complicated given the widespread nature of 2-PP-induced CSPs in COQ8A. Lack of CSPs in the N-lobe rules out this domain as the 2-PP binding site. If it is assumed that 2-PP does not bind in the nucleotide binding site (assumption addressed below), then the most probable location would be somewhere at the interface of the KxGQ domain and C lobe. It is plausible that binding in this location could have an allosteric effect on chemical shifts in the KxGQ domain and the β7/β8-αE/αF motif in the C-lobe.

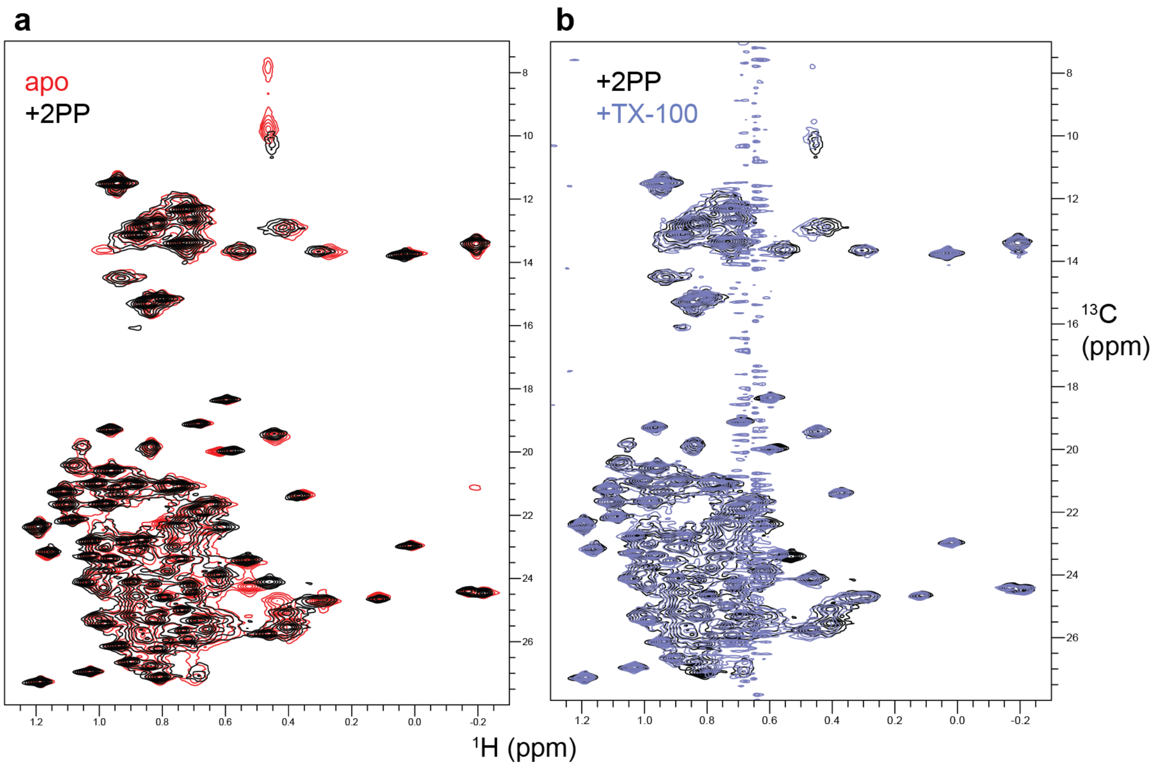


**Figure 1.** 2-PP and TX-100 induce chemical shift perturbations in COQ8A. (**a**) $^1H$-$^{13}C$ HMQC spectra of COQ8A in the apo state (red) and with 1 mM 2-PP. (**b**) $^1H$-$^{13}C$ HMQC spectra of COQ8A with 1 mM 2-PP (same sample as in **a**) before (black) and after (blue) addition of 1 mM TX-100. The streak in **b** is due to the methyl group in TX-100.

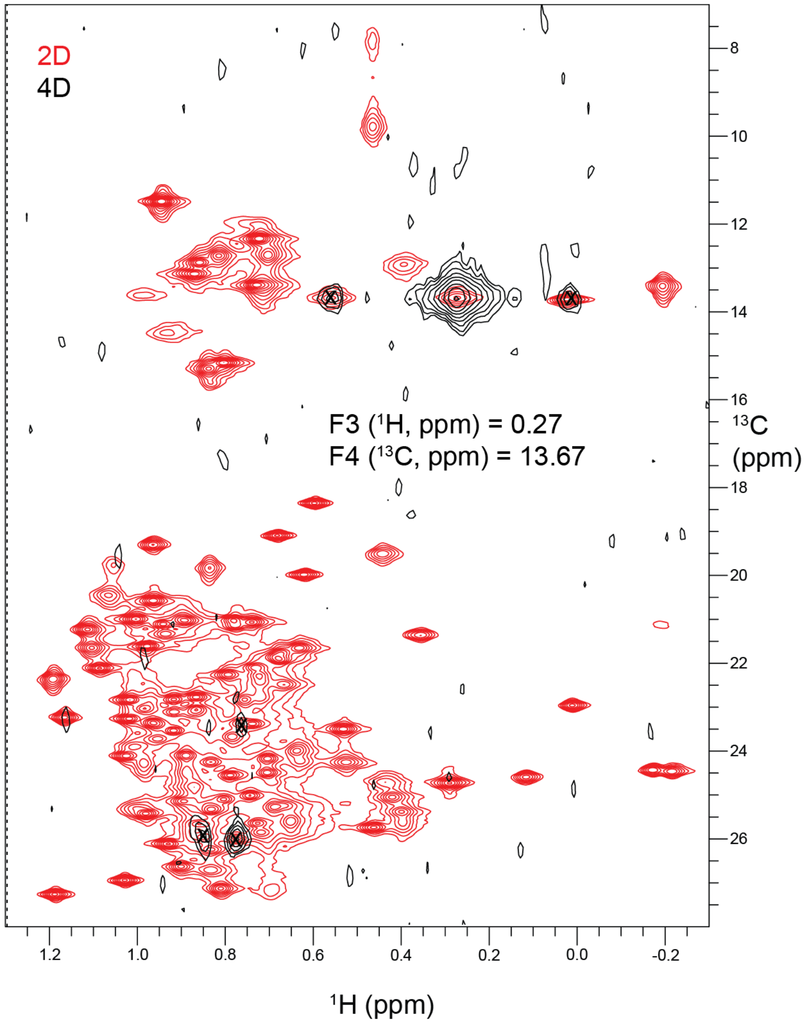


**Figure 2.** Example 2D plane from 4D NOESY spectrum. The 2D HMQC spectrum of COQ8A is shown in red, while the 2D plane corresponding to the specified F3 and F4 frequencies in the 4D NOESY are in black. The intense peak at the 0.27, 13.67 ppm corresponds to the 'diagonal peak' in a 3D spectrum. Other peaks in this plane, marked with an x, are NOEs.

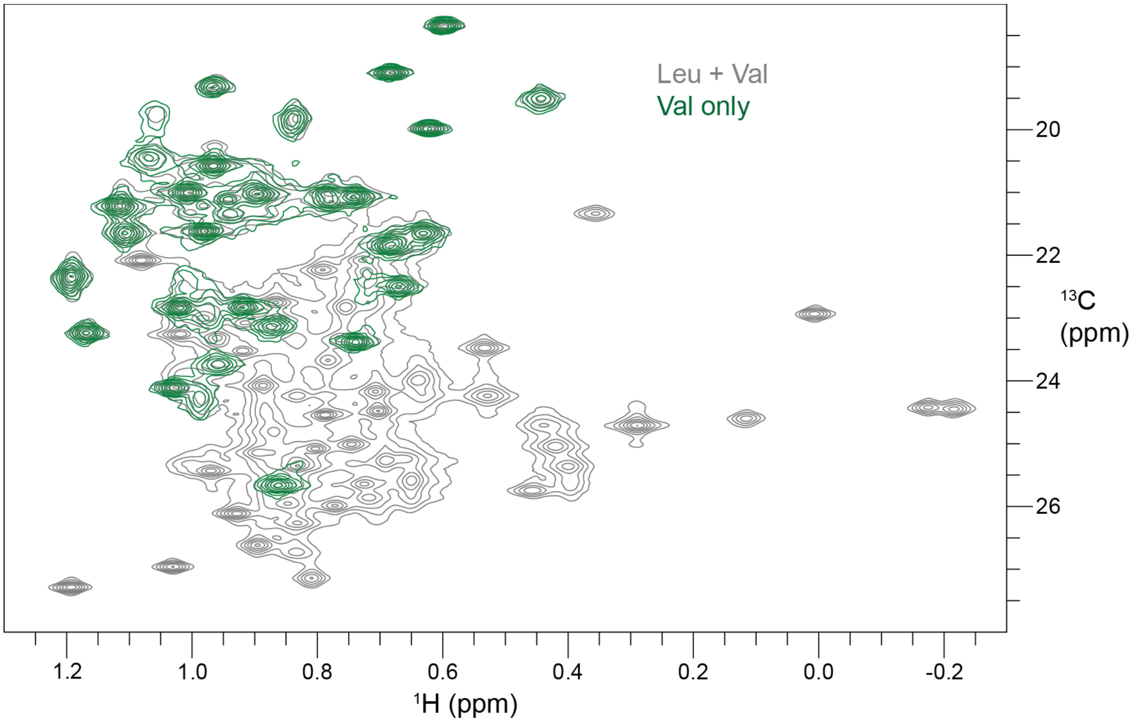


**Figure 3.** Val-selective labeling allows for L/V residue type discrimination. Overlay of $^{1}$H-$^{13}$C HMQC spectra of apo COQ8A where both L and V residues are labeled (grey) and where only V is labeled (green).

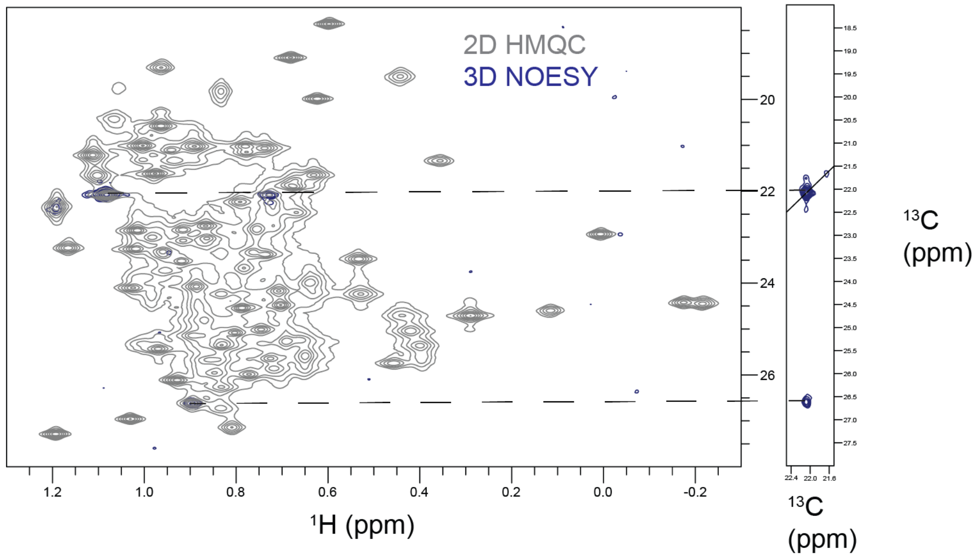


**Figure 4.** Dimethyl labeling of L/V residues allows pairing of geminal methyl groups. HC (left, blue) plane 3D NOESY experiment is overlaid with the 2D HMQC (grey) on the right. The corresponding CC plane is shown on the right. The off-diagonal peak corresponds to the geminal methyl group.

#### References


1. Ollerenshaw, J. E., Tugarinov, V. & Kay, L. E. Methyl TROSY: explanation and experimental verification. *Magn Reson Chem* 41, 843–852 (2003).

2. Gorman, S. D., Sahu, D., O'Rourke, K. F. & Boehr, D. D. Assigning methyl resonances for protein solution-state NMR studies. *Methods San Diego Calif* 148, 88–99 (2018).

3. Tugarinov, V., Kay, L. E., Ibraghimov, I. & Orekhov, V. Y. High-resolution four-dimensional 1H-13C NOE spectroscopy using methyl-TROSY, sparse data acquisition, and multidimensional decomposition. *J Am Chem Soc* 127, 2767–75 (2005).

4. Pritišanac, I. *et al.* Automatic Assignment of Methyl-NMR Spectra of Supramolecular Machines Using Graph Theory. *J Am Chem Soc* 139, 9523–9533 (2017).

5. Nerli, S., Paula, V. S. D., McShan, A. C. & Sgourakis, N. G. Backbone-independent NMR resonance assignments of methyl probes in large proteins. *Nat Commun* 12, 691 (2021).
