## Supplementary material for "Small Molecule Modulation of the Archetypal UbiB protein COQ8": Synthesis Supplement

### Supplementary Synthesis Information

##### TPP-UNC-CA157 Synthesis

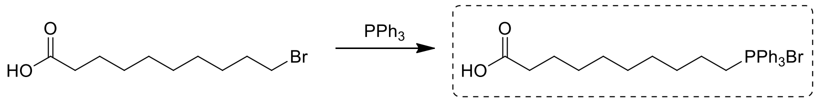


**50530-12-6** **2**

A mixture of 10-bromodecanoic acid (500 mg, 1.99 mmol, 1.00 eq) and $PPh_3$ (522 mg, 1.99 mmol, 1.00 eq) in MeCN (5.00 mL) was was stirred at 90 °C for 70 h. LC-MS showed desired MS peak was detected. The mixture was concentrated to give crude compound (**2**) (1.10 g, 73.2% yield) as a light pink oil, and used directly for next step without further purification.

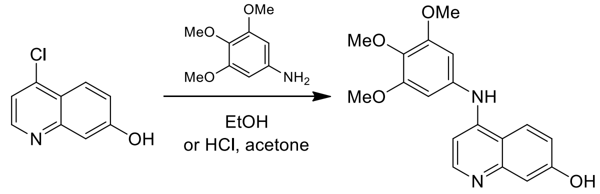


**181950-57-2** **1**

To a solution of 4-chloroquinolin-7-ol (500 mg, 2.78 mmol, 1.00 eq) and 3,4,5-trimethoxyaniline (2.04 g, 11.1 mmol, 4.00 eq) in acetone (30.0 mL) was added HCl (12.0 M, 1.00 mL, 4.31 eq). The mixture was stirred at 75 °C for 16 h. LC-MS showed desired MS peak was obvious. The reaction was cooled to room temperature, poured into ice water, sodium hydroxide solution to adjusted pH to 9 by adding aqueous NaOH (2M), extracted with ethyl acetate (100 mL*3), the organic phase was dried over $Na_2SO_4$, filtered and concentrated to dryness under reduced pressure. The residue was purified by flash silica gel chromatography (ISCO®; 12.0 g Sepa Flash® Silica Flash Column, Eluent of 0~75% Ethyl acetate/Petroleum ether gradient @ 45 mL/min) to afford Compound **1** (700 mg, 2.12 mmol, 76.3% yield) as a light yellow solid.

**$^1$H NMR:** (400 MHz, MeOD)

δ: 8.00-8.07 (m, 2H). 6.95-7.02 (m, 1H), 6.80-6.85 (m, 1H), 6.63-6.69 (m, 2H), 6.56-6.62 (m, 1H), 3.81 (s, 6H), 3.76 (s, 3H).

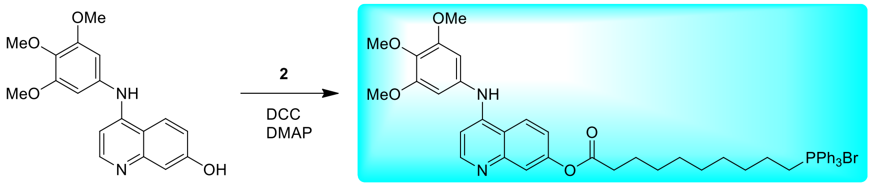


**1**

**Target 3**
**25 mg, 95.0%**

A mixture of compound **(1)** (200 mg, 613 umol, 1.00 eq) , compound **(2)** (315 mg, 613 umol, 1.00 eq) and DMAP (3.74 mg, 30.6 umol, 0.05 eq) and N,N'-dicyclohexylmethanediimine (126 mg, 613 umol, 124 uL, 1.00 eq) in $CH_2Cl_2$ (10 mL) was was stirred at 20 °C for 24 h under $N_2$ atmosphere. LC-MS showed desired compound was detected. The mixture was concentrated under reduced pressure to afford crude. The crude was purified by chromatography column ($SiO_2$, EtOAc/Methanol = 20:1~5:1), and then prep-HPLC (column: Phenomenexluna 30*30mm*10um+YMC AQ 100*30*10um; mobile phase: [water (0.1%TFA)-ACN]; B%: 15%-65%, 26 min) to afford compound **Target 3** (150 mg, 157 umol, 25.6% yield) as a yellow oil.

**$^{1}$H NMR:** (400 MHz, $CDCl_3$)

δ: 11.27 (s, 1H), 8.96 (d, $J$ = 9.26 Hz, 1H), 8.19 (d, $J$ = 4.75 Hz, 1 H), 7.77-7.86 (m, 4 H), 7.57-7.74 (m, 12H), 7.27-7.34 (m, 1H), 6.81 (d, $J$ = 6.50 Hz, 1H), 6.68 (s, 2H), 3.87 (s, 3H), 3.82 (s, 5H), 3.13 - 3.30 (m, 2H), 2.54 (t, $J$ = 6.82 Hz, 2H), 1.68 (m, 4H), 1.55 (d, $J$ = 6.00 Hz, 2H), 1.19-1.45 (m, 10H).
